# Strain-specific alterations in gut microbiome and host immune responses elicited by *Bifidobacterium pseudolongum*

**DOI:** 10.1101/2022.03.16.484607

**Authors:** Bing Ma, Samuel J. Gavzy, Vikas Saxena, Yang Song, Wenji Piao, Hnin Wai Lwin, Ram Lakhan, Jegan Lyyathurai, Lushen Li, Michael France, Christina Paluskievicz, Marina WillsonShirkey, Lauren Hittle, Arshi Munawwar, Emmanuel F. Mongodin, Jonathan S. Bromberg

**Author notes:** Division of Lung Diseases, National Heart, Lung, and Blood Institute (NHLBI), National Institutes of Health (NIH), Bethesda, Maryland, USA. Corresponding authors: Bing Ma; Jonathan S. Bromberg.

## Abstract

The beneficial effects attributed to *Bifidobacterium* are thought to arise from their host immunomodulatory capabilities, which are likely to be species- and even strain-specific. However, their strain-specificity in direct and indirect immune modulation remain largely uncharacterized. We have shown that *B. pseudolongum* UMB-MBP-01, a murine isolate, is capable of suppressing inflammation and reducing fibrosis *in vivo*. To ascertain the mechanism driving this activity and to determine if it is specific to UMB-MBP-01, we compared it to *B. pseudolongum* type strain ATCC25526 of porcine origin using a combination of *in vitro* and *in vivo* experimentation and comparative genomics approaches. Despite many shared features, we demonstrate that these two strains possess distinct genetic repertoires in carbohydrate assimilation, differential activation signatures and cytokine responses in innate immune cells, and differential effects on lymph node morphology with unique local and systemic leukocyte distribution. Importantly, the administration of each *B. pseudolongum* strain resulted in major divergence in the structure, composition, and function of gut microbiota. This was accompanied by markedly different changes in intestinal transcriptional activities, suggesting strain-specific modulation of the endogenous gut microbiota as a key to host responses of immune modulation and changes in intestinal *B. pseudolongum* strains. These observations highlight the importance of strain-specificity characteristics of *Bifidobacterium* for prophylactic supplementation for immune modulation and advance our understanding of the mechanisms which drive the association between *Bifidobacterium* and health benefit.

## Introduction

*Bifidobacterium* spp. are naturally occurring residents within the gastrointestinal (GI) tract of mammals and are typically considered beneficial [1, 2]. Due to their purported health-promoting properties, *Bifidobacterium* spp. have been incorporated into many live biotherapeutic (LBP) prophylactic formulations, mostly known for applications in alleviating intestinal inflammatory conditions [3–7]. The potential mechanisms underlying the health benefits of *Bifidobacterium* include the suppression of growth of gut pathogens [8, 9], capabilities to alter gut metabolism and to enhance epithelial barrier function [10, 11], and anti-inflammatory modulation of host immunity [12–15]. In particular, their immunomodulatory properties are not limited to the direct effects on GI tissues, but also indirect effects enacted through their influence on the gut microbiota [16]. *Bifidobacterium* spp. are known to participate in mutualistic interactions with endogenous intestinal microorganisms that can subsequently evoke both immediate as well as delayed immune responses [17, 18]. However, the cellular and molecular underpinnings *Bifidobacterium*’s biotherapeutic effects remain unclear with contradictory findings reported [19]. Fundamentally important questions such as what specific mechanisms through which they exert immunomodulatory effects, to what extent the interactions with the gut microorganisms affect the immune responses, and what are the roles of elicited intestinal responses in these processes remain outstanding.

The immunomodulatory properties of individual *Bifidobacterium* spp. are strain-dependent, despite similar effects produced by closely related strains (i.e., alleviation of lactose intolerance or improved host antimicrobial activity) [20–22]. In fact, immunomodulatory effects are independent of microbial phylogeny [20]. Recent investigations suggested that differences in cell wall composition and structure might be responsible for strain-specific immunomodulatory effects [23]. Microorganism-associated molecular patterns (MAMPs) possess variable biochemistry, even between strains, serving as microbial stimuli that orchestrate molecular cascades in the host immune response and mucosal homeostasis [24–26]. Exopolysaccharide (EPS) and pili may play a role in *Bifidobacterium’s* strain-specific pro-homeostatic immunomodulation [27, 28]. Other molecular mechanisms such as lipoteichoic acid and specific metabolites such as acetate could also contribute to strain-specific immunity [26, 29, 30]. Comparisons of the immunomodulatory properties of closely related strains can be leveraged to identify which are strain-specific and to characterize the microbial determinants of specific host responses, which will provide the basis to rationally hone biotherapeutics for prophylactic applications [15, 31, 32].

We previously showed, using a major histocompatibility complex (MHC)-mismatched murine cardiac transplant model, that fecal microbiota transfer (FMT) caused shifts in the gut microbiota which profoundly influenced allograft outcomes [33]. FMT using stool samples from healthy pregnant mice (immune suppressed) resulted in improved long-term allograft survival and prevented inflammation and fibrosis in grafts, as compared to FMT using stool samples from colitic or nonpregnant control mice [33]. *B. pseudolongum* was revealed as a microbial biomarker for the pregnant mouse gut microbiota, from which we subsequently isolated and sequenced UMB-MBP-01 [34]. Importantly, gavage with UMB-MBP-01 alone reproduced the same improved graft outcomes as FMT using whole stool of pregnant mice, implicating this strain as one of the main responsible microbes [33]. Thus, the murine tropic strain UMB-MBP-01 may serve as a model organism to investigate the mechanisms of microbe-driven immunomodulation.

In this study, we performed a genome-wide comparison of UMB-MBP-01 to all other *B. pseudolongum* genomes, including three additional *B. pseudolongum* strains (E, EM10, EM13) isolated from the same feces sample of a pregnant mouse, as well as to the porcine tropic type strain ATCC25526, in order to investigate the genetic attributes underlying the immunomodulatory properties. Further, we revealed distinct effects on local and systemic immunity induced by UMB-MBP-01 and ATCC25526, using both *in vitro* and *in vivo* approaches. Importantly, the oral administration of the two *B. pseudolongum* strains resulted in profound alterations in composition, structure and function of the murine gut microbiome, accompanied with markedly different intestinal transcriptome activities. These observations suggest that modulation of the endogenous gut microbiome is a key element of *Bifidobacterium* immunomodulatory attributes. A deeper understanding of the strain specificity and mechanisms of action through which specific strains regulate host responses will facilitate the clinical translation of live therapeutics and the development of potential immunomodulatory therapy targets.

## Results

### High genome plasticity of *B. pseudolongum* reflects strong host adaptability

The pangenome of *B. pseudolongum* was constructed using 79 strains including the 4 strains sequenced as part of this study (**Supplemental Table 1A**). Homologous gene clusters (HGCs) were identified in this set of genomes based on all-versus-all sequence similarity (**Supplemental Table 1B**). A total of 4,321 *B. pseudolongum* HGCs were revealed, among which 31.7% were core (present in almost all strains), 57.0% were dispensable (singleton or present in very few genomes), and the remaining 11.3% were considered accessory. *B. pseudolongum* demonstrated a smaller pangenome size that was 87.8% of *B. breve* and 59.5% of *B. longum* pangenomes (**Supplemental Figure 1**). *B. pseudolongum* had the fewest number of conserved HGCs (N=1,370) but the largest proportion of dispensable pangenome (57.0%) compared to the two other *Bifidobacterium* species *B. longum* and *B. breve* that were both human-associated. This disproportionally large dispensable pangenome may be indicative of strong niche adaptation capabilities of *B. pseudolongum*, reflecting its broad host range, being widely distributed among mammals [35].

Whole genome sequencing was performed on three *B. pseudolongum* strains (E, EM10, and EM13) isolated from the same pregnant mice feces as UMB-MBP-01 (sequencing statistics in **Supplemental Table 2A**). Comparison among the four murine strains revealed 1,520 shared coding DNA sequence (CDS), which comprised 97.2% of UMB-MBP-01 coding genes (**Supplemental Table 1C**). 107 CDS were conserved in at least two but not in all four genomes, and 37 CDS were strain-specific. Most of these genes had unknown functions, and those with known functions related to bacteriophage assembly and function (i.e., capsid protein, integrase, transposes, bacteriophage replication gene, cell lysis protein, microvirus H protein) or carbohydrate hydrolysis and transport (glycosyl hydrolases, ABC transporter permease). On the other hand, comparison between UMB-MBP-01 and ATCC25526 revealed 1,351 shared CDS (86.4% of UMB-MBP-01 coding genes), and 157 genes that belonged to one strain but not the other (**Supplemental Table 1D**). Interestingly, most of these strain-specific genes also belonged to the categories of bacteriophage assembly and functions as well as carbohydrate hydrolysis and transport, in addition to genes with unknown function.

Together these data suggested bacteriophage-mediated transduction was a major contributor to dissemination of carbohydrate metabolism capabilities, potentially through horizontal gene transfer among closely related murine-derived strains, as well as more distantly related *B. pseudolongum* strains.

Whole genome Average Nucleotide Identity (ANI) clustering suggested two subspecies, *B. pseudolongum* subsp. *pseudolongum* clade that contained ATCC25526, and *B. pseudolongum* subsp. *globosum* clusters that had three distinct clades I-III (**Figure 1**). Subspecies *globosum* clade III had the largest number of coding genes (1,642+/-70) among all clades and contained UMB-MBP-01 and the three isolates from the source stools of pregnant mice. The subspecies *pseudolongum* clade had the smallest number of coding genes among all clades (1,519+/-35.6). Overall, 1,599 HGCs accounted for 37.0% of *B. pseudolongum* pangenome were identified as clade-specific (>90% genes belonging to the same clade), and the majority originated from *globosum* clade III (N=648), while clade *pseudolongum* provided the fewest (N=130). The large number of clade-specific genes found in *globosum* clade III genomes suggested a high degree of genome plasticity to facilitate adaptation to cope with environmental heterogeneity. Further functional enrichment analyses revealed *globosum* cluster III-specific HGCs were mostly involved in periplasmic transport systems, permeases and glycoside hydrolases (GHs), particularly the families GH29 (α-L-fucosidase), GH3 (β-glucosidase) and GH31 (α-glucosidase) (**Supplemental Table 1E**). No GH families were enriched in any of the other clades. Together, UMB-MBP-01 and ATCC25526 belonged to two different subspecies, each of which comprises considerable genetic variation. The genome of UMB-MBP-01 contained more clade-specific genes and was enriched for genetic features in carbohydrate metabolism to assimilate greater varieties of glycans, presumably facilitating its niche adaptive capabilities in the glycan-rich gut environment.

**Figure 1.**
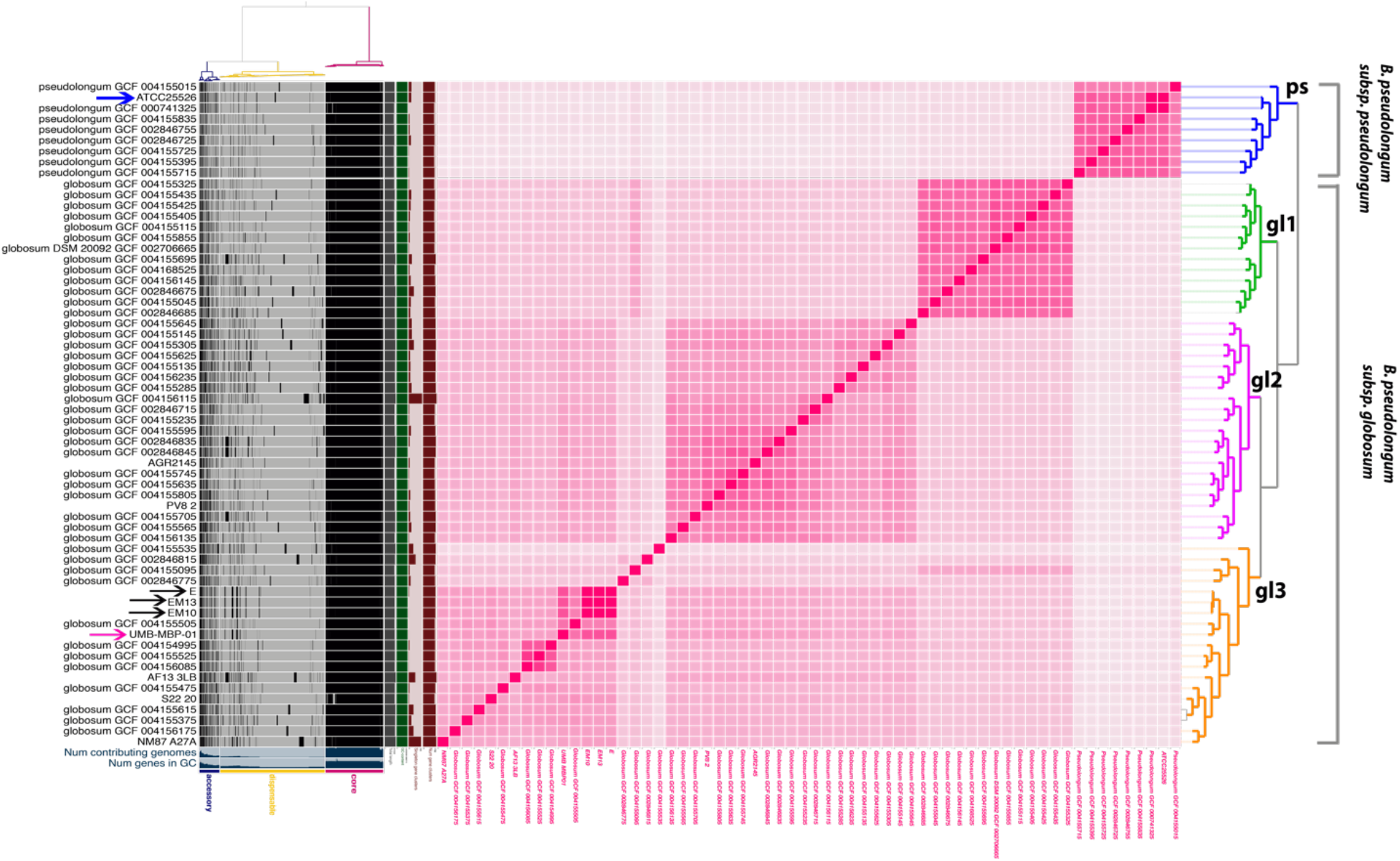
Pangenome analyses of *B. pseudolongum* genomes. Pangenome constructed using 79 strains, including the 5 strains sequenced in this study (**Supplemental Table 3**) and displayed using anvi’o *vers* 6.2 [69]. Homologous gene clusters (HGCs) were identified based on all-versus-all sequence similarity in left panel and categorized as core, accessory or dispensable depending on their level of conservation. Genome ANI (Average Nucleotide Identity) was calculated using Sourmash vers 3.3 [71]. Blue arrows indicate the two strains compared: ATCC25526 and UMB-MBP-001. Black arrows indicate the other three *B. pseudolongum* strains isolated from the source stool of pregnant mice.

### Specialized carbohydrate metabolizing capabilities of UMB-MBP-01 and ATCC25526

The abundance of *Bifidobacterium* glycolytic features is reflective of their metabolic adaptation to the complex carbohydrate-rich GI tract [36, 37]. We performed *in silico* prediction of the carbohydrate fermentation capabilities to comprehensively investigate glycan-assimilation capabilities for all 79 *B. pseudolongum* genomes, using with the Carbohydrate-Active enZYmes Database (CAZy) database [38]. This analysis revealed 236 genes of *B. pseudolongum* pangenome encoding predicted carbohydrate-active enzymes from 34 glycosyl hydrolase families, 14 glycosyl transferase families and eight carbohydrate esterase families (**Supplemental Table 4A**). Only 33.5% of the carbohydrate-active enzyme coding genes belonged to the core pangenome. Core GHs included those mostly responsible for the breakdown of plant-derived polysaccharides (*i.e.*, starch) and a wide range of other carbohydrates, such as GH13 (glycosidase), GH77 (α-amylase), GH43 (β-xylosidase), GH36 (α-galactosidase), GH2 (β-galactosidase), GH3, and GH6 (cellobiohydrolases). Notably, GH13 is the enzyme family known to be most commonly found in *Bifidobacterium* genomes and active on a wide range of carbohydrates including the plant-derived starch and the related substrates of trehalose, stachyose, raffinose, and melibiose [37, 39]. Conversely, 47.9% of the identified carbohydrate-active enzymes genes were found in the dispensable pangenome. The *globosum* clade II (N=72) and III (N=58) encoded most of these enzymes, while the *pseudolongum* clade encoded the least (N=13). These results demonstrated the highly specialized carbohydrate assimilation gene repertoires of different strains, particularly in *globosum* clade II and III.

Using UMB-MBP-01 as the reference for all other *B. pseudolongum* strains, both conserved and specific glycohydrolases capabilities were revealed (**Supplemental Figure 2, Supplemental Table 1F, 4B**). Interestingly, the clusters based on GH are mostly in agreement with the clades generated based on ANI, suggesting distinct carbohydrates assimilation capabilities of different *B. pseudolongum* clades. GH29, GH31, GH42 (β-galactosidase), and ABC-type polysaccharide transport permease genes were most prevalent in *globosum* clade III that contained UMB-MBP-01. Further, GH36, GH2, and GH94 (cellobiose phosphorylase) were found absent in subspecies *pseudolongum* clade but prevalent in *globosum* clade III. In particular, an uncommon GH23 family (peptidoglycan lyses) was only observed in UMB-MBP-01 and the three other isolates from the pregnant mouse. Overall UMB-MBP-01 and ATCC25526 share some enzymatic capabilities in metabolizing dietary polysaccharides and host-derived glycogens, while also having specialized glycohydrolases genes.

We further characterize the carbohydrate utilization capabilities of UMB-MBP-01 and ATCC25526 using anaerobic microplates pre-coated with various carbon sources. Out of the 95 carbon sources tested, the two strains demonstrated the same capabilities on 86 (90.5%) (**Supplemental Table 3**), including key carbon sources N-acetyl-D-glucosamine, D-fructose, L-fucose, α-D-glucose, glucose-6-phosphate, maltose, maltotriose, D-mannose, D-sorbitol, and pyruvic acid. Two relatively uncommon sugars D-melibiose and D-raffinose could be metabolized by ATCC25526 but not UMB-MBP-01. On the other hand, D-galactose, D-gluconic acid, D-glucosaminic acid, glycerol, D-mannitol, α-ketovaleric acid, and D, L-lactic acid were uniquely metabolized by UMB-MBP-01. This result is in principle in an agreement of the specific GH families predicated *in silico*. Together these data indicated a wide range of carbohydrate metabolizing capabilities ranging from dietary to host-derived glycans for both strains, while UMB-MBP-01 had specialized capabilities to metabolize galacto-oligosaccharides.

We sought to characterize the secretome of *B. pseudolongum* by examining protein localization based on the presence of a signal peptide [40]. Proteins which are secreted extracellularly have the potential to directly interact with the other gut microorganisms and with host tissues [27, 41] (**Supplemental Table 5A**). Overall, the sec-dependent secretion machinery, but not the twin-arginine (Tat) system, was conserved in all *B. pseudolongum* strains, indicating protein translocation function was conserved but likely occurs only in the unfolded state [42]. Secreted proteins were more likely to be part of the dispensable genome (73% of secreted proteins versus 53% of cytoplasmic proteins; **Supplemental Table 5B**), indicating a high degree of diversity in the secretome among strains of *B. pseudolongum*. Proteins which were predicted to be extracellularly secreted include solute-binding proteins of ABC transporter systems, amidases related to the peptidoglycan hydrolysis, glycosyl hydrolyses, cell surface proteins that make up pilus subunits, and cell wall-degrading peptidases. Interestingly, the secretome of the clade containing ATCC25526 was enriched for collagen adhesion proteins (**Supplemental Table 5C**) but lacked multiple secreted GH25 extracellular proteins. These proteins are prevalent in the clade which includes UMB-MBP-01 and are involved in the binding and hydrolysis of peptidoglycan (**Supplemental Table 5D**). As peptidoglycan components were implicated in important aspects of mucosal immunological signaling [43], this may contribute to varied immunomodulatory capabilities between UMB-MBP-01 and ATCC25526.

Differential activation and cytokine responses in dendritic cells and macrophages induced by *B. pseudolongum* strains ATCC25526 and UMB-MBP-01

To understand the immunomodulatory impact of the two *B. pseudolongum* strains, bone marrow derived dendritic cells (BMDC) and peritoneal macrophages (MΦ) were treated with UV-killed whole bacteria (cells) or isolated *Bifidobacterium* exopolysaccharide (EPS). We first examined the effect of these treatments on expression of costimulatory receptors. For BDMCs, treatment with either *B. pseudolongum* strain stimulated increased CD40 and CD86 expression, although CD86 expression was significantly greater after treatment by ATCC25526 than UMB-MBP-01 (P < 0.01) (Figure 2A-D). Treatment with ATCC25526 cells stimulated increased MHC class II, while treatment with UMB-MBP-01 cells stimulated increased CD80 expression. Neither UMB-MBP-01 EPS nor ATCC25526 EPS altered expression of these surface receptors on BMDCs. For MΦ, ATCC25526 cells stimulated increased CD40, while ATCC25526 EPS did not (**Figure 2E-G**). The other cell surface receptors were not affected by treatment with bacterial cells or EPS for MΦ. Overall, ATCC25526 and UMB-MBP-01 bacterial cells, but not EPS, triggered activation of important costimulatory receptors on innate myeloid cells *in vitro*, and these bacterial strains differed in these activities.

**Figure 2.**
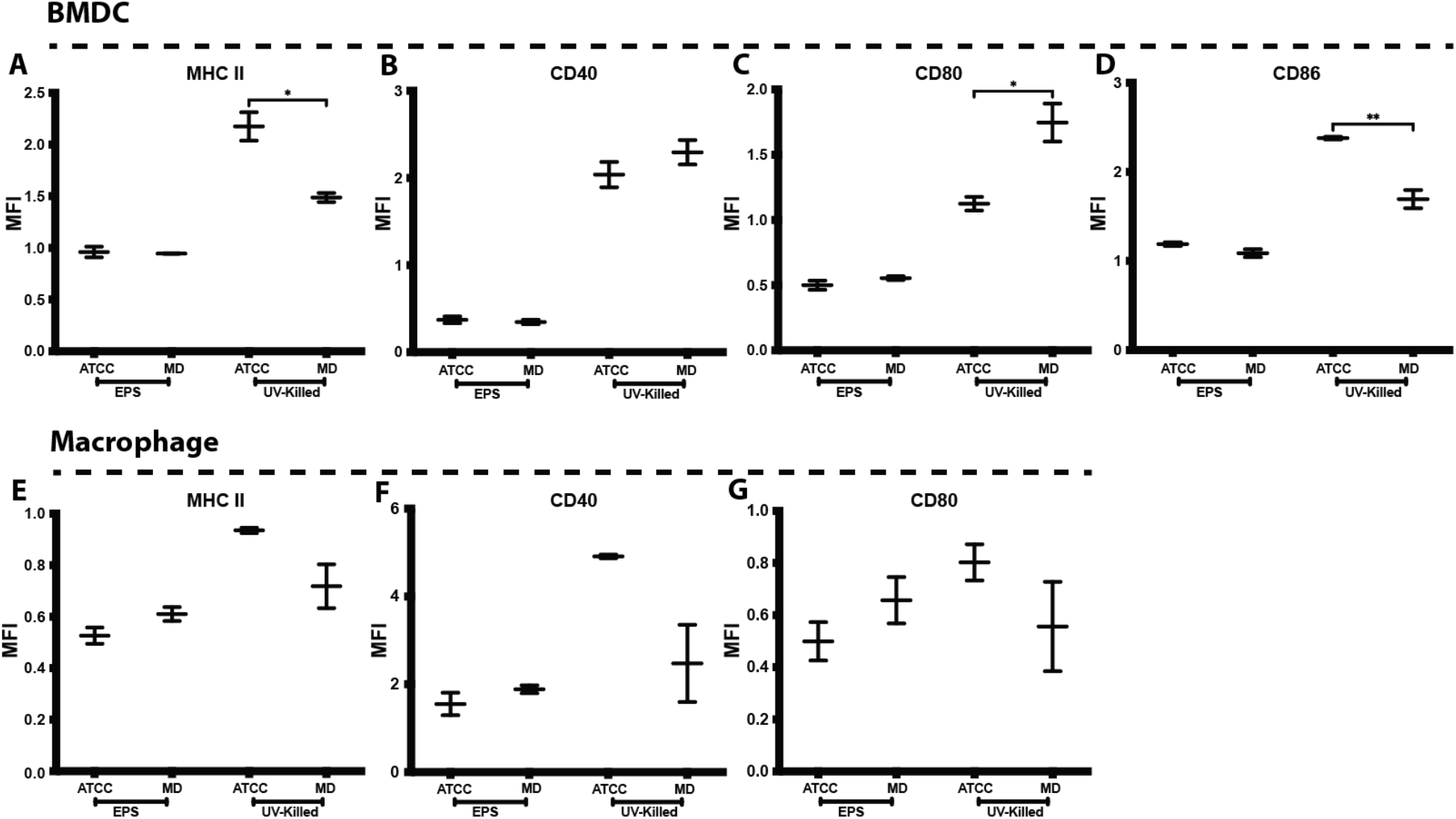
*Bifidobacterium* alters DC and MΦ surface phenotype. DC and MΦ cultured with media alone, ATCC25526 (ATCC) or UMB-MBP-01 (MD) UV-killed *Bifidobacterium* or EPS derived from each strain. After 24 hrs of culture, cells analyzed by flow cytometry. DC gated on live CD11c+, and MΦ gated on live F4/80+ populations. DC stained for **A)** MHC class II, **B)** CD40, **C)** CD80, and **D)** CD86. MΦ stained for **E)** MHC class II, **F)** CD40 and **G)** CD80. MFI: mean fluorescence intensity; DC: Bone marrow derived dendritic cells; MΦ: peritoneal macrophages; MFI values normalized to control and compared using one-way ANOVA. * p value < 0.05; ** p value < 0.01. MΦ data representative of two experiments. DC data represented as merge of one experiment with EPS and one experiment with UV killed bacteria, each data set is normalized to its respective control.

We next examined the effect of *Bifidobacterium* and EPS alone on cytokine production in innate immune cells. Both BMDC and MΦ showed increased secretion of IL-6, TNF*α*, and IL-10 when stimulated with UMB-MBP-01 and ATCC25526 strains. Induction of cytokine expression was also strain-specific as ATCC25526 cells stimulated a greater increase in IL-6 and IL-10 than UMB-MBP-01 cells in BMDCs (**Figure 3A, 3C**); and TNF*α* expression was also increased to a greater extent by

**Figure 3.**
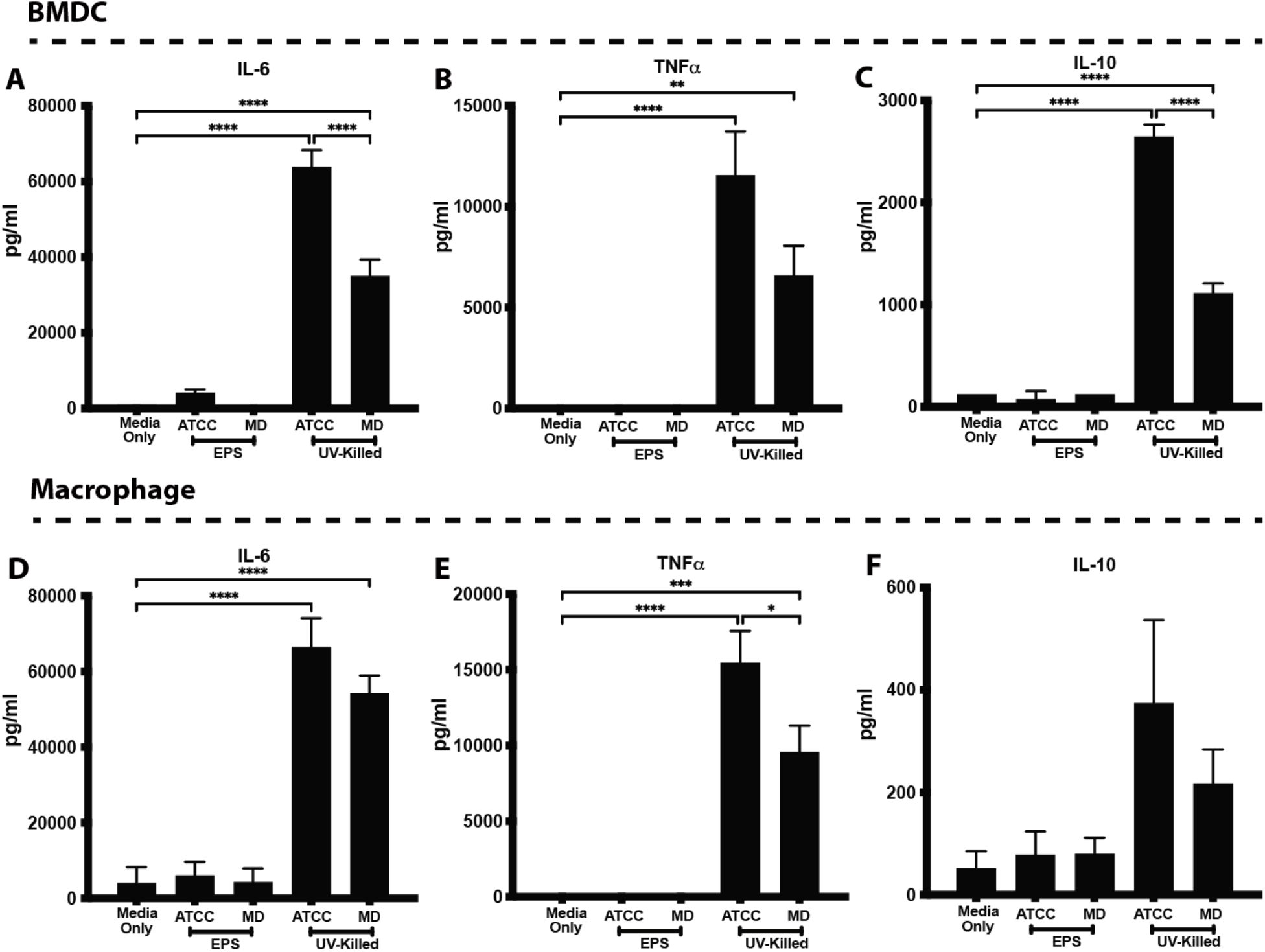
*Bifidobacterium* alters DC and MΦ cytokine secretion. DCs **(A-C)** or MΦ **(D-F)** stimulated with EPS or UV-killed ATCC25526 (ATCC) or UMB-MBP-01 (MD), and 24 hrs later supernatants analyzed for **A, D)** IL-6, **B, E)** TNF*α*, and **C, F)** IL-10 by ELISA. Treatments compared using one-way ANOVA. * p value < 0.05; ** p value < 0.01, *** p value < 0.001, **** p value < 0.0001. Data representative of two separate experiments.

ATCC25526 compared to UMB-MBP-01 with a borderline statistical significance (p = 0.059, **Figure 3B**). For MΦ, treatment with ATCC25526 cells increased TNF*α* compared to UMB-MBP-01 (**Figure 3E**), whereas there were no strain-specific differences in IL-6 or IL-10 (**Figure 3D, 3F**). *B. pseudolongum* EPS did not stimulate cytokine production in either BMDC or MΦ. Similar to activation of co-stimulatory receptors, UMB-MBP-01 and ATCC25526 cells elicited unique myeloid cell cytokine responses that differed from one another and were not recapitulated by EPS.

### *Bifidobacterium* strains induce distinct changes in local and systemic leukocyte distribution and lymph node morphology

We next assessed whether *Bifidobacterium* strains differentially induced changes in immune cell distribution and lymph node (LN) architecture *in vivo* using a mouse model. As illustrated in **Figure 4A**, mice received broad spectrum antibiotics for 6 days, a regimen that depleted endogenous microbiota [44], followed by oral gavage with each bacterial strain or their EPS, and then daily immunosuppression with tacrolimus (3 mg/kg/d s.c.). Two days after gavage, mesenteric and peripheral LNs (MLN and PLN) and intestinal tissues were harvested, and the intraluminal fecal content was collected for gut microbiome characterization. The effect of these microbiota on the distribution of immune cell populations was assessed by immunohistochemistry of intestinal segments and flow cytometry and immunohistochemistry of LNs.

**Figure 4.**
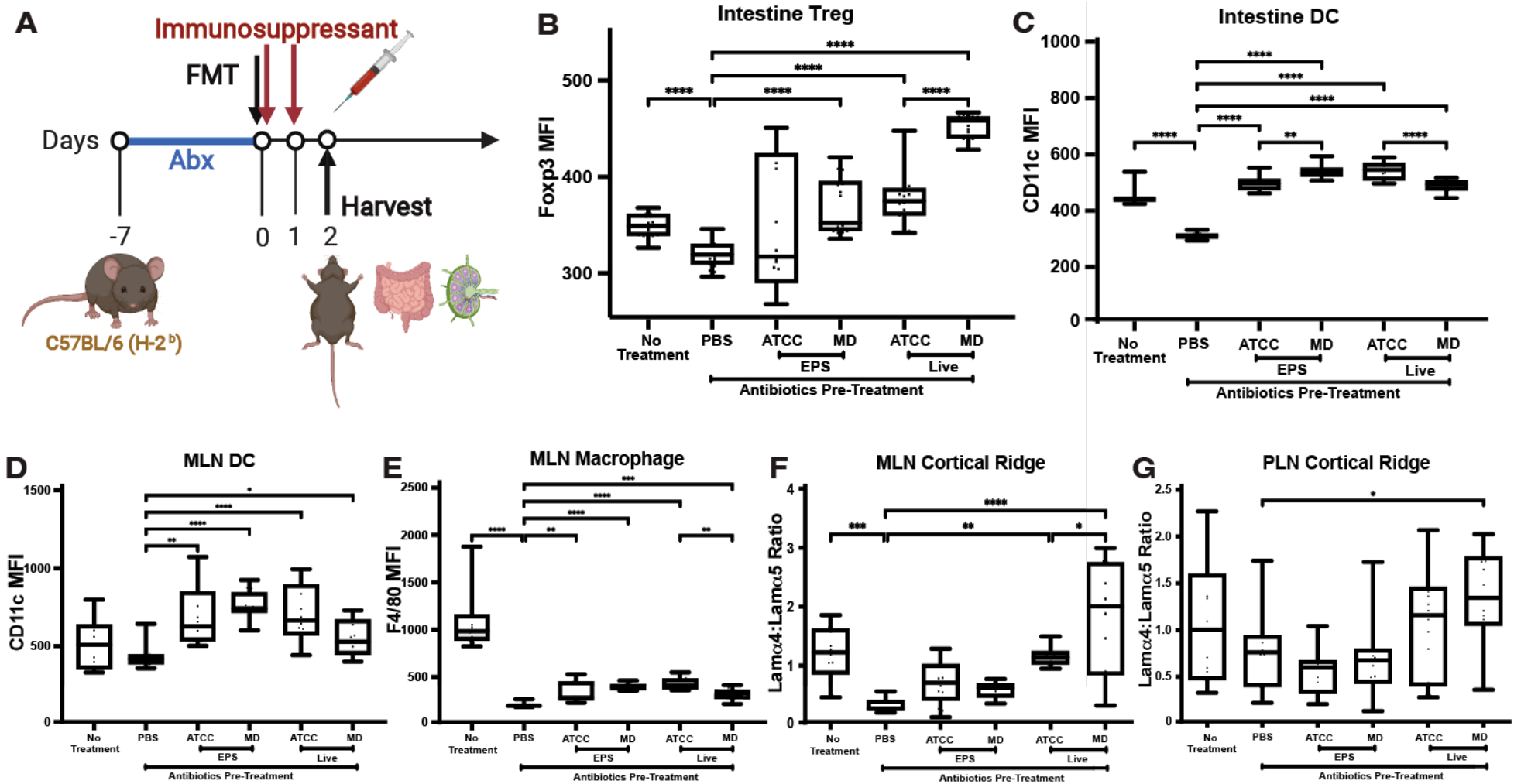
*Bifidobacterium* strains induce unique changes in local and systemic immune cell distribution and LN architecture. **A)** Experimental design. C57BL/6 mice treated with antibiotics for 6 days followed by gavage with *B. pseudolongum* ATCC25526 (ATCC), UMB-MBP-01 (MD), or PBS (control). Mice then treated with the immunosuppressant tacrolimus for the next two days. Tissues harvested 2 days after FMT. Frozen sections of small intestine stained for **B)** Treg (anti-Foxp3 mAb), and **C)** DC (anti-CD11c mAb). MLN sections stained for **D)** DC (anti-CD11c mAb) and **E)** MΦ (anti-F4/80 mAb). LN stained for laminin α4 and laminin α5 and laminin α4:α5 ratio calculated for **F)** MLN CR, and **G)** PLN CR. MFI: mean fluorescence intensity. MLN: mesenteric lymph nodes; PLN: peripheral lymph node. MFI values normalized using the sum of mean, and categories compared using one-way ANOVA. * p value < 0.05; ** p value < 0.01, *** p value < 0.001, **** p value < 0.0001. Data representative of two separate experiments, 3 mice/group.

Within the small intestine, both UMB-MBP-01 and ATCC25526 cells resulted in significantly more Foxp3+ regulatory T cells (Treg) compared to PBS control, while UMB-MBP-01 resulted in significantly more Treg compared to ATCC25526 (**Figure 4B**, flow cytometry gating protocol listed in **Supplemental Figure 3**). Gavage with UMB-MBP-01 EPS, but not ATCC25526 EPS, resulted in increased Treg compared to control (**Figure 4B**). ATCC25526 and UMB-MBP-01 cells resulted in more DC compared to PBS control, while ATCC25526 cells also resulted in more DC compared to UMB-MBP-01 cells (**Figure 4C).** UMB-MBP-01 and ATCC25526 EPS resulted in increased DC compared to PBS control, while UMB-MBP-01 EPS also resulted in more DC compared to ATCC25526 EPS (**Figure 4C**). Intestinal MΦ did not significantly change after whole bacteria or EPS gavage.

By flow cytometry, we observed decreased MLN Treg in UMB-MBP-01 EPS treated animals, but otherwise there were no other differences in the number and proportion of innate myeloid (DC, MΦ) or adaptive lymphoid (CD4 T cells, CD8 T cells, Treg, and B cells) cells in the MLN or PLN of mice treated with the *B. pseudolongum* strains compared to those treated with antibiotics alone or untreated controls (**Supplemental Figure 4A-L)**.

For LN immunohistochemistry, we focused on the LN cortex because our previous work showed that architectural and cellular changes within this zone were most critical in mediating immune tolerance and suppression [45]. Within the MLN and PLN T cell cortex, neither gavage with whole bacteria nor EPS affected the number of Treg present, as measured by immunohistochemistry. This contrasts with the flow cytometry results above which showed that UMB-MBP-01 EPS caused a decrease in MLN Treg (**Supplemental Figure 4D)**. The difference between the flow cytometry and histologic results for Treg is likely due to the focus of histology analysis only on cells in the cortex while the flow cytometry summates all the cells in the entire LN. Gavage with whole bacteria and EPS from both strains increased DCs in the cortex of MLN compared to control (**Figure 4D**), while the number of DC enumerated by flow cytometry did not change in MLN or PLN (**Supplemental Figure 4 E, K)**, with these differences again likely due to the factors noted above. MΦ increased in the cortex of MLN after whole bacteria or EPS treatments from both strains (**Figure 4E**), but not in PLN (**Supplemental Figure 4N**). ATCC25526 cells also resulted in a greater increase in MLN MΦ compared to UMB-MBP-01 cells (**Figure 4E**). This result contrasts with flow cytometry data where there was no difference in MΦ populations in MLN or PLN after treatment.

Overall, gavage with whole bacteria and EPS altered gut associated innate myeloid cells and Tregs without affecting systemic distribution, as evidenced by unchanged PLN populations (**Supplemental Figure 4 M, N)**. Gavage with whole *Bifidobacterium* cells led to increased intestinal Treg and DCs as well as increased MLN MΦ and DCs compared to controls. EPS also produced most of these effects apart from ATCC25526 EPS lack of effect on intestinal Treg compared to control. UMB-MBP-01 cells led to increased intestinal Treg compared to ATCC25526 cells. In contrast, ATCC25526 cells led to increased DC in both intestine as well as increased cortical MLN MΦ compared to UMB-MBP-01 cells. UMB-MBP-01 EPS caused increased intestinal DC compared to ATCC25526 EPS. In contrast to the *in vitro* findings where EPS was generally inactive, *in vivo* treatment with EPS alone stimulated similar innate myeloid cell and Treg increases in gut and MLN compared to increases induced by live bacterial cells.

Since LN stromal fiber structures are important mediators of immune responses [46], we next assessed LN architecture using the ratio of laminin α4 to laminin α5 in the LN T cell cortex of the cortical ridge (CR) and around the high endothelial venules (HEV). An increased laminin α4:α5 ratio is indicative of immune tolerance and suppression [45]. In the MLN CR, both UMB-MBP-01 and ATCC25526 increased the laminin α4:α5 ratio, with UMB-MBP-01 cells causing a greater increase compared to ATCC25526 cells (**Figure 4F**). The laminin α4:α5 ratio was not changed around the MLN HEV by either whole bacteria or EPS **(Supplemental Figure 4O)**. In PLN CR, only UMB-MBP-01 cells resulted in an increased laminin α4:α5 ratio (**Figure 4G**). Overall, gavage with both *B. pseudolongum* strains increased local MLN CR laminin α4:α5 ratios, while only UMB-MBP-01 increased the laminin α4:α5 ratio in systemic PLN CR. This effect was more prominent for UMB-MBP-01 compared to ATCC25526, demonstrating strain-specific differences in immune modulation.

### Markedly different intestinal transcriptional activities in response to UMB-MBP-01 than to ATCC25526

To determine the effect of UMB-MBP-01 and ATCC25526 on host gene expression, we characterized the transcriptome of mouse intestinal tissues harvested two days after gavage with either UMB-MBP-01, ATCC25526, or no bacteria control. Differentially expressed genes (DEGs) were identified by comparing the two treatment groups to the control and revealed both shared and strain-specific effects on transcription (**Supplemental Table 6B-D**). A total of 420 and 425 DEGs were observed in comparisons of UMB-MBP-01 vs. control and ATCC25526 vs. control, respectively, and 139 DEGs were observed comparing UMB-MBP-01 to ATCC25526 directly. Based on the log_2_ fold change (LFC) scale of DEGs, the strongest intestinal response was elicited by UMB-MBP-01, compared to either ATCC25526 or control (**Supplemental Figure 5**). Functional enrichment analyses revealed the effects elicited by UMB-MBP-01 were mainly involved in positive regulation of cell activation, leukocyte and lymphocyte activation, B cell activation, and somatic recombination of immunoglobulin superfamily domains (**Supplemental Figure 5A, C**). Elicited effects of ATCC25526 include responses in phagocytosis, membrane invagination, defense response to bacterium and complement activation (**Supplemental Figure 5E**). These results further supported our observations that ATCC25526 elicited distinct host responses compared to UMB-MBP-01, which induced greater numbers of DEGs and stronger host responses.

We further examined the host responses present in both UMB-MBP-01 and ATCC25526 as well as those present only in one but not the other, to pinpoint the differential host responses induced by the two strains. Of the DEGs identified comparing UMB-MBP-01 or ATCC25526 to the control (n=411 and 416), 59.6% and 58.9%, respectively, were identified in both comparisons (**Figure 2A, Supplemental Table 6**). These overlapped DEGs (N=238) and the condition-specific DEGs, that included 164 DEGs only up-regulated in UMB-MBP-01 versus control and 111 DEGs only upregulated in UMB-MBP-01 versus ATCC25526, comprised the majority of all DEGs (85.9%). Downregulated genes accounted for only a small fraction of all DEGs (14.1%) and majority of them were identified in only the comparison of UMB-MBP-01 vs.

ATCC25526 (N=67, 80.1% of downregulated). The DEGs that were upregulated in both UMB-MBP-01 vs. control and ATCC25526 vs. control include B cell immunity, collagen metabolism, immunoglobulin protein expression, cytokines (IL-1*β*, IL-10, IL-13, IL-21), TNF receptor superfamily, among others (**Supplemental Table 6**). Two functional pathways were enriched in UMB-MBP-01 vs control, but not in ATCC25526 vs control: regulation of cell-cell adhesion and T cell activation and the response to interferon γ and interferon β (**Figure 5B)**. In contrast, the host responses to ATCC25526 but not UMB-MBP-01 were enriched in functions involved in fatty acid metabolism, lipid localization, acylglycerol metabolism, and cholesterol and sterol homeostasis (**Supplemental Figure 5D**). Together these data further indicated that the effects of UMB-MBP-01 or ATCC25526 were mediated through different pathways. The ATCC25526 strain appeared to exert immunomodulatory effects, at least in part, via stimulation of phagocytosis and induced lipid metabolism, while the UMB-MBP-01 strain exerted stronger effects, mostly through upregulating antibody secretion and regulation of multiple aspects of lymphocyte functions, including cytokines, adhesion, and activation.

**Figure 5.**
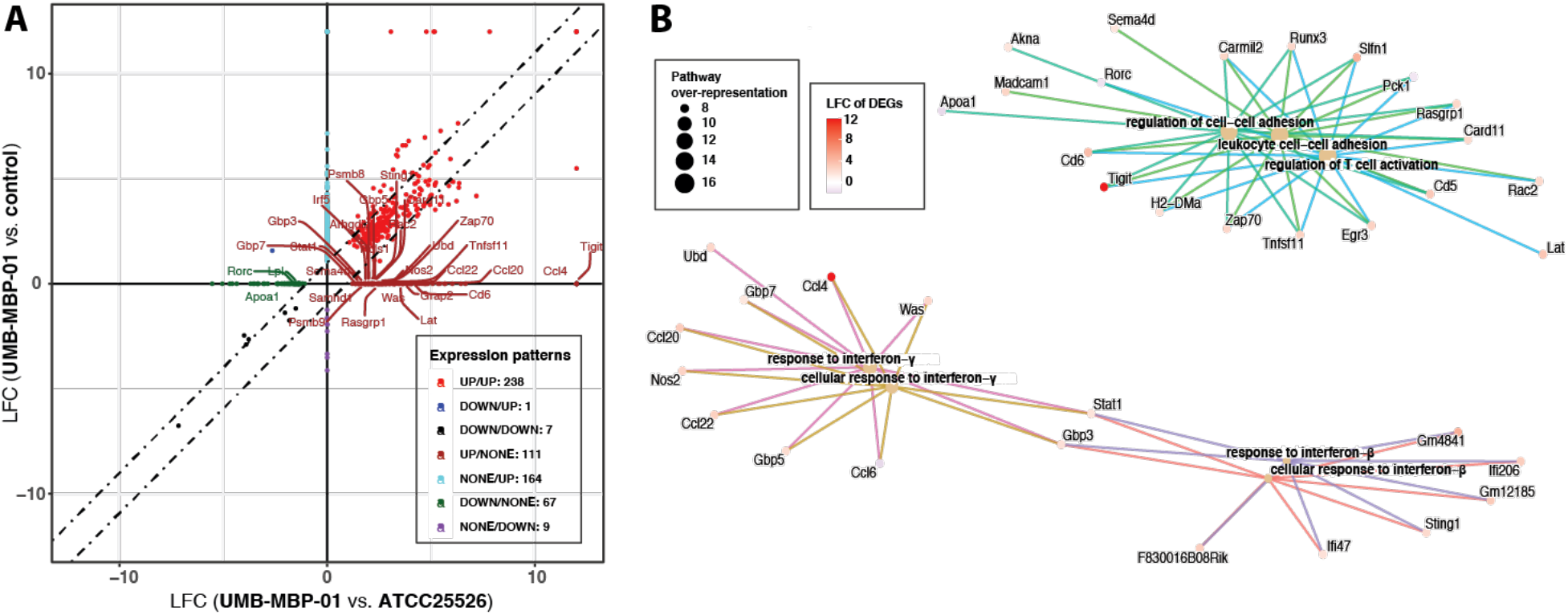
Transcriptome profiling of intestinal tissues in response to ATCC25526 or UMB-MBP-01. **A**) Quadrant plot to show whether differential expressed genes (DEGs) have the same or opposite relationships between each of the pairwise comparison of UMB-MBP-001 vs control and UMB-MBP-001 vs ATCC25526. DEGs were determined using log2 fold change (LFC) >(+/-)1 and false discovery rate (FDR)<0.05. **B**) Gene-Concept network for most over-represented Gene Ontology (GO) terms to depict over-represented functions based on q-value and gene-count. Over-representation analyses [99] of DEGs that are only different abundant in UMB-MBP-001 vs control but not in ATCC25526 vs control, using GO ontologies performed using enrichGO function of clusterProfile Bioconductor package[100]. For pairwise comparison enrichment analyses for any two conditions, please refer to **Supplemental Figure 4**.

**Figure 6.**
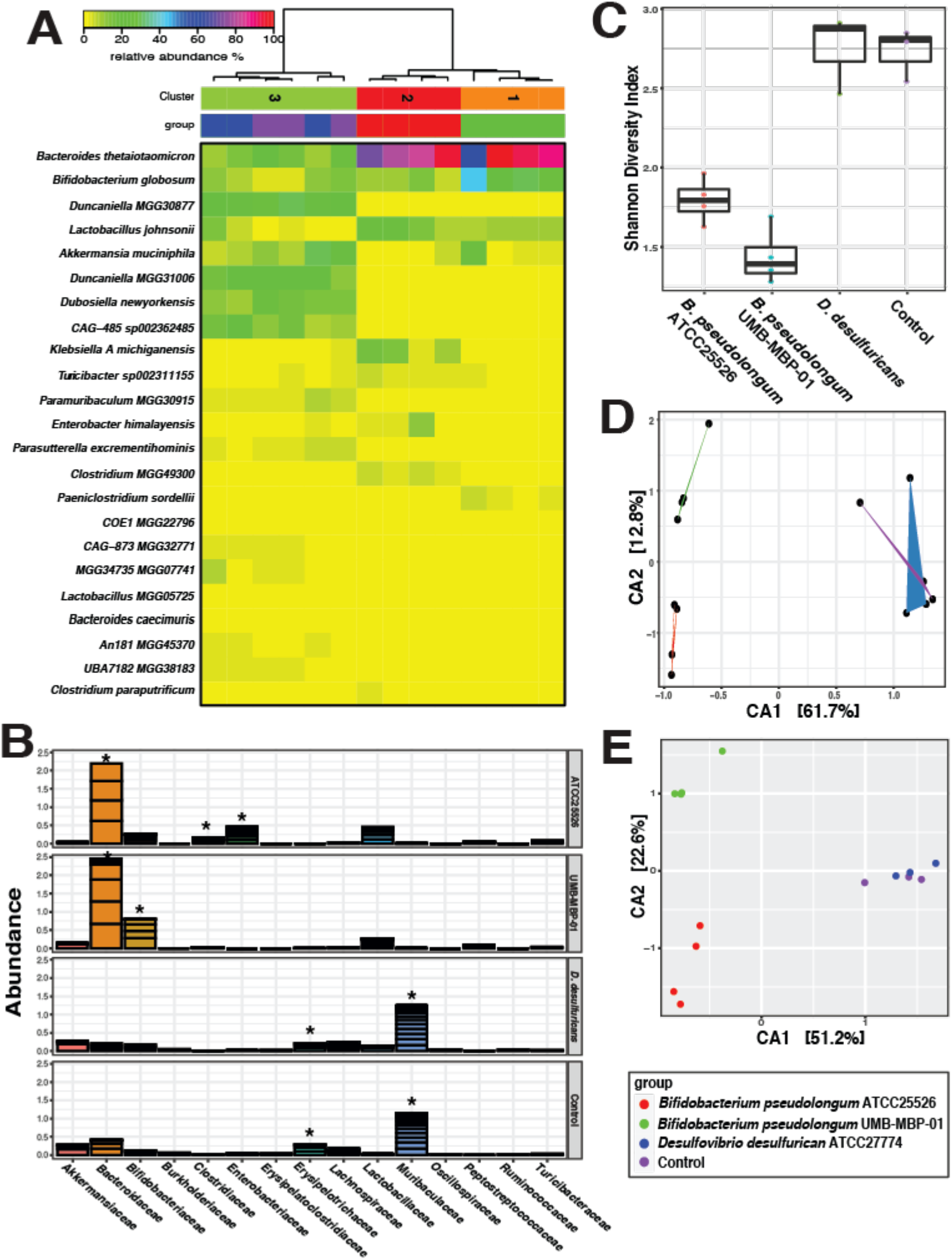
Alterations in gut microbiome after bacterial engraftment. **A**) Heatmap of the top 20 most abundant intestinal bacterial taxa relative abundance in mice intraluminal samples. Ward linkage clustering based on Jensen-Shannon distance was calculated using the vegan package in R [94]. Taxonomic profiles of the microbial community were characterized using the comprehensive mouse gut metagenome (CMGM) catalog[47]. **B**) Cumulative abundance of major bacterial families. The relative abundances of each family are stacked in order from greatest to least, and are separated by a horizontal line. **C**) Shannon diversity index (within-community diversity) of the four experimental groups. **D**), **E**) Canonical Correspondence Analysis (CCA) of microbial functional pathways characterized using HUMAnN2 (v0.11.2)[91] and Uniref90 database [90] based on Bray-Curtis distance. CA1 and CA2 selected as the major components based on the eigenvalue.

As extracellular molecules may play an important role in eliciting immunomodulatory effects, we also compared intestinal gene expression following gavage with live *B. pseudolongum* bacteria to that with *B. pseudolongum* produced EPS. The comparison revealed DEGs that were mostly group-specific without much overlap with EPS vs. control group (12.1%, N=39) (**Supplemental Figure 6A, B**). These data indicated that the predominant intestinal transcriptional responses were due to *B. pseudolongum* whole cell gavage (62.6%, N=201), compared to DEGs in EPS gavage (25.5%, N=82). Gene-pathway network analyses indicated B cell receptor activation and signaling, antigen-receptor mediated signaling, and phagocytosis recognition and engulfment were highly upregulated by whole *B. pseudolongum gavage* (**Supplemental Figure 6C**). While both live cells and EPS induced antimicrobial circulating immunoglobulin, the transcriptional effects were an order of magnitude higher for live cells (**Supplemental Figure 6C-D**). This result was commensurate with the observations above that EPS did not stimulate cytokine production or cell surface costimulatory receptor expression in either BMDC or MΦ. Together, our data suggested that the immunomodulatory effects of *B. pseudolongum* were potentially via their metabolic activities and secreted molecules and/or other cell membrane components other than the surface structure of EPS.

Different *B. pseudolongum* strains elicit rapid, profound alterations in both structure and function of gut microbiome

We next investigated the impact of bacterial gavage on the gut microbiome using shotgun metagenomic sequencing of the intraluminal fecal content (40.6±7.7 million reads per sample; **Supplemental Table 2B**). Taxonomic composition was established using the comprehensive mouse gut metagenome catalog (CMGM) [47] designed specifically to characterize the mouse gut microbiome (**Figure 5A, Supplemental Table 7**). A significant reduction in gut microbial community diversity was observed after both UMB-MBP-01 and ATCC25526 gavage, with UMB-MBP-01 gavage resulting in the lowest diversity (**Figure 5C**). After *B. pseudolongum* administration, the most outstanding changes in specific taxonomic groups were the marked increases in the relative abundance of *Bacteroides thetaiotaomicron* and *Lactobacillus johnsonii* and relative depletion of *Muribaculaceae* and *Erysipelotrichaceae* (**Figure 5B, Supplemental Figure 7**). Gavage with the *Desulfovibrio* did not produce a significant change in the microbiome, and *Muribaculaceae*, *Erysipelotrichaceae*, and *Lachnospiraceae* were the most abundant groups in these communities and in the not-treatment control communities. These data indicated the gavage of either *B. pseudolongum* strains profoundly altered the gut microbial community, with a significant reduction in abundance of endogenous gut microorganisms.

Canonical Correspondence Analysis (CCA) on both community taxonomic profiles and functional pathways resulted in concordant clustering patterns, in that ATCC25526 or UMB-MBP-01 each resulted in a distinct community, and that were clearly separate from *Desulfovibrio*-treated and no bacteria controls (**Figure 5D, 5E**). Based on linear discriminant analysis (LDA) effect size (LEfSe) analysis [48], UMB-MBP-01 resulted in a significantly higher abundance of *B. pseudolongum* than ATCC25526 (19.8±6.1% vs. 6.1%±2.0%, P = 0.02, **Supplemental Figure 8A,B**). These data suggested murine strain UMB-MBP-01 was better able to colonize the mouse gut than the porcine isolate ATCC25526, lead to greater abundance in the murine gut environment. On the other hand, *Enterobacteriaceae* (*Klebsiella michiganensis* and *Enterobacter himalayensis*), and *Clostridiaceae* (*Clostridium paraputrificum* and *Clostridium* MGG49300) were significantly more enriched in the ATCC25526 treated group but were mostly absent in the UMB-MBP-01 gavage mice (**Supplemental Figure 8C-F**). Based the scaled eigenvalue, the top taxa and pathways that contributed to the separation of the clusters in ordination analyses were identified (**Supplemental Figure 9A, B, Supplemental Table 8**). The ATCC25526 cluster was attributed to *Enterobacteriaceae* and *Clostridiaceae*, and the top contributors included *K. pneumonia*, *K. michiganensis*, *B. animalis*, *E. himalayensis*, and *Clostridium paraputrificum*. The most prominent pathways attributed to ATCC25526 cluster include motility (peptidoglycan maturation), gluconeogenesis, energy conversion (fatty acid β-oxidation), and L-threonine biosynthesis. On the other hand, *Akkermansia muciniphila*, *Paeniclostridium sordellii*, and *B. pseudolongum* were among the top significant contributors to the UMB-MBP-01 cluster. The most outstanding pathways for UMB-MBP-01 included ribonucleotide and amino acid biosynthesis (folate transformation, L-isoleucine, L-arginine, L-lysine) and pyruvate fermentation (pyruvate/acetyl-CoA pathway). Together, the data indicated UMB-MBP-01 or ATCC25526 each altered the gut microbiota profoundly and distinctively, which may contribute to their distinct immunomodulatory effect.

## Discussion

*B. pseudolongum* demonstrates great intraspecies genetic diversity and shows patterns consistent with host specificity, rendering it an advantageous model organism to study the effect of intraspecies variation on host immunomodulation [35]. Further, as a predominant species in the murine GI tract, *B. pseudolongum* displays an extensive enzymatic capacity and might act as a keystone species in this environment [35, 49]. In this study, we employed the murine strain UMB-MBP-01, which demonstrates an anti-inflammatory and pro-homeostatic effect [33, 34], and porcine-isolated *B. pseudolongum* type strain ATCC25526 to investigate the strain-specific mechanisms of host responses *in vitro* and *in vivo*. The distinct genetic attributes and immunomodulatory capabilities of UMB-MBP-01 from the *B. pseudolongum* type strain ATCC25526 show that *Bifidobacterium* modulates intestinal responses and host immunity in a strain specific manner. We observed UMB-MBP-01 exerted stronger immunologic effects in intestinal responses mostly likely through regulation of multiple aspects of lymphocyte functions, while ATCC25526 appeared to exert immunomodulatory effects, at least in part, via stimulation of phagocytosis and induced lipid metabolism. We further demonstrated the *in vitro* strain-specific activation and cytokine responses in DC and MΦ and changes in local and systemic leukocyte distribution and LN morphology, demonstrating the unique immune modulatory effects of the two *B. pseudolongum* strains. A deeper understanding of strain-specific immunomodulatory properties is fundamentally important to inform probiotic design as well as immunomodulatory therapeutic targeting.

It remains unclear whether *B. pseudolongum* immune and intestinal modulation is mediated through direct interactions with the intestinal epithelium, or indirectly via modulation of endogenous gut microbiome with consequent effects on intestinal metabolism and immunity, or both [18]. In our study, the administration of two separate *B. pseudolongum* strains resulted in profoundly different gut microbiomes in both structure and functional capabilities as well as intestinal responses, suggesting the critical involvement of the endogenous gut microbiome as a key element of their immunomodulatory attributes and indicating that indirect effects likely contribute. Our results align with recent key clinical findings, suggesting that *Bifidobacterium* could act as a “microbiome modulator” to competitively exclude toxigenic pathogens and orchestrate homeostatic gut metabolism and host immune responses [50–53]. It is worth noting that the effect of LBPs on the structure and function of the microbial community has not been accepted among standard parameters to characterize or evaluate LBP efficacy. Our study emphasizes the importance of characterization of the dynamics of the gut microbial community to understand the efficacy and specificity of LBPs.

Distinctive from ATCC25526 and pro-inflammatory bacterial control, UMB-MBP-01 demonstrated strain persistence within the gut microbiome after administration, further highlighting the role of gut microbiome in LBP-host interaction. Previous studies suggested that the administration of *B. longum* subsp*. longum* has stable, persistent colonization in recipients whose gut microbiome previously had low abundance of gene content involved in carbohydrate utilization, suggesting competition for resources as a key mechanism determining strain persistence [54]. This may relate to the ecological concept of “colonization resistance’, whereby endogenous microbiota occupy host tissues with an intrinsic capability to limit the introduction of exogenous microorganisms and the expansion of endogenous microorganisms, while a microbiota with low capacity in carbohydrate assimilation could be more permissive to exogenous colonization that can fundamentally disrupt the microbiota [55, 56]. Given human gut reliance on microbiota to cope with glycan-rich gut environment for the metabolism of luminal oligosaccharides [57], the ability of *Bifidobacterium* strains to utilize complex carbohydrates provides a selective advantage to effectively compete for nutrients with other bacteria in the gut microenvironment [57]. Interestingly, their repertoire of glycoside hydrolases are species- or strain-specific [39, 58], indicating indispensable roles for glycan-assimilation in their specific niche adaptability to the intestinal microenvironment. Supporting the competitive exclusion ecological theory, we observed the depletion of endogenous gut microorganisms *Muribaculaceae* and *Erysipelotrichaceae* with *Bifidobacterium* treatment, as well as the specialized carbohydrate metabolism of UMB-MBP-01 in utilizing a greater variety of oligosaccharide molecules and host-derived glycans. Future longitudinal characterization of strain persistence, microbiota changes, and oligosaccharide and glycoprotein assimilation may hold a key to determine probiotic strain specificity in host adaption and intestinal responses.

The bacterial determinants of immunomodulation properties of individual strains remains an underdeveloped research area. Recent studies revealed strains of the same species induced different immunophenotypes, suggesting that bacterial-induced immunomodulation is not dictated by bacterial phylogeny [22, 59]. Multiple LBP produced effector molecules that interact with host immunity have been recently identified [60]. In particular, cell wall components found in Gram-positive bacteria, such as peptidoglycan and lipoteichoic acid, contain MAMPs which are recognized by immunoregulatory pattern recognition receptors such as Toll-like receptors [23].

*Bifidobacterium* peptidoglycans have also demonstrated immunomodulatory effects on the Th1 polarization of naïve T cells as well as DC maturation and enhanced immune responses [61, 62]. Other surface molecules such as EPS and pili also play a role in *Bifidobacterium’s* strain-specific pro-homeostatic immunomodulation [27, 28]. We also observed here expanded enzymatic capabilities of UMB-MBP-01, but not ATCC25526, in assimilating oligosaccharide molecules and host-derived glycans as well as the capacity to manufacture cell wall components such as peptidoglycan. Together these studies implicate the genetic variation of different bacterial strains as underlying induced intestinal responses and mucosal immunological signaling. This speculation warrants direct experimental validation.

Making conclusions about the immunomodulatory effects of bacterial strains based on the surface structures alone is inaccurate, as the preparations of the cell wall components and extracellular polysaccharides are strongly influenced by cultivation conditions [63]. The differences observed in this study between *Bifidobacterium* whole cells and purified EPS *in vitro* vs. *in vivo* indeed suggest the presence of many other molecules and mechanisms that contribute to the regulation of immunity and inflammation. In particular, the observation that EPS alone yielded gut Treg recruitment in addition to innate myeloid cell populations suggests that EPS may act indirectly through an intermediary, such as intestinal epithelial cells, which then influence leukocyte subsets. Indeed, characterization of *B*. *pseudolongum*-induced innate immune responses revealed that these bacteria induce a more balanced anti-inflammatory and homeostatic cytokine response from DC and MΦ [64]. In addition, *B. pseudolongum* induced changes in LN architecture, resulting in an increased ratio of extracellular matrix protein laminin α4 to laminin α5 in the cortical ridge, a microdomain structure that is mechanistically associated with immunologic suppression and tolerance [33]. These observations demonstrate that a single probiotic bacterial strain can influence local, regional, and systemic immunity through both innate and adaptive pathways. A holistic understanding of the strain-specific bacterial effects is critical to inform probiotic design as well as immunomodulatory therapeutic targeting.

## Conclusion

The distinct genetic attributes and immunomodulatory capabilities of UMB-MBP-01 compared to *B. pseudolongum* type strain ATCC25526 show that *Bifidobacterium* modulates intestinal responses and host immunity in a strain-specific manner. Our results highlight the importance to characterize individual *Bifidobacterium* strains and not to generalize their immunomodulatory effects to other strains of the same species, despite their many shared features. It is critical to investigate both endogenous microbiota in response to LBP strains and to profile the intestinal responses, in order to interrogate mechanistically the highly coordinated multicellular host-microbe interactions, which are key to understanding strain-specific immunomodulation. Future studies are warranted to investigate the specific bioactive metabolites and pathways through which the gut microbiota exert their immunomodulatory effects, and the specific intestinal cell types that respond to those signals. A comprehensive understanding of strain-specific immunomodulatory properties is fundamentally important to inform probiotic design as well as immunomodulatory therapeutic targeting.

## Methods and Materials

### Strains cultivation and genomic sequencing

*B. pseudolongum* strain ATCC25526 was purchased from ATCC (Manassas, Virginia). *B. pseudolongum* strain UMB-MBP-01 was isolated from the feces of C57BL/6J mice through passages and screening on Bifidus Selective Medium (BSM) agar (Sigma-Aldrich, St. Louis, MO, USA), as previously described [34]. Both strains were initially grown anaerobically at 37°C for 3–5 days on Bifido Selective Media (BSM) agar plates (Millipore Sigma, Burlington, MA), from which a single colony was selected and grown in BSM broth (Millipore Sigma, 90273-500G-F) until stationary phase (up to 3 days). *Desulfovibrio desulfuricans subsp. desulfuricans* (ATCC27774) was purchased from ATCC and grown in ATCC Medium: 1249 Modified Baar’s Medium (MBM) for sulfate reducers, which was made according to ATCC protocol. Cultures were initially incubated under anaerobic conditions for 5 days on Modified Baar’s Medium agar plates, after which single colonies were chosen, transferred to liquid media, and incubated for up to 3 weeks. *B. pseudolongum* strains ATCC25526 and UMB-MBP-01 were used in cell stimulation and cytokine assays.

*B. pseudolongum* UMB-MBP-01 was sequenced previously [34]. For strain ATCC25526, genomic DNA extraction was performed using a lysozyme/mutanolysin-based cell lysis followed by purification using the Wizard Genomic DNA Purification Kit (Promega, Madison, WI, USA). Library preparation on extracted DNA was conducted using a Kapa kit (Roche, Indianapolis, IN) for 150-bp paired-end sequencing, and sequencing was performed with an Illumina (San Diego, CA) MiSeq system. Sequencing was performed by the University of Maryland School of Medicine, Institute for Genome Sciences, Genomics Resource Center with standard operating procedures and assembled using SPAdes v3.14.0 [65]. Contig ordering was performed using MAUVE contig mover [66] and the UMB-MBP-01 genome as reference [67].

### Exopolysaccharide (EPS) Isolation

EPS was extracted from strains X and Y using the protocol previously published by Bajpai and colleagues [68]. Briefly, bacterial cultures were grown in 500 ml BSM media to early stationary phase, after which trichloroacetic acid was added (14% v/v final) and the mixture incubated at 37°C for 40 minutes. After centrifugation at 8,000 g for 20 minutes at 4°C, the supernatant was collected, absolute ethanol added (2:1 v/v EtOH:sup), followed by incubation at 4°C for 48h and centrifugation at 8,000xg at 4°C for 20 minutes. This ethanol wash step was repeated to remove any impurities, and a final centrifugation at 8,000xg at 4°C for 20 minutes was performed. The resulting pellet was then dissolved in 5 to 10 ml of water, before being dialyzed against DI water for 48 hours, and lyophilized.

### Anaerobic microplate assay

Anaerobic microplates (AN plates, Biolog, Hayward, CA) pre-coated with 95 various carbon sources was used in the assay. Each well of the 96-well AN Biolog plate was coated by a sole-carbon source, with one well being used as no carbon control. Metabolism of the substrate in particular wells results in formazan production, producing a color change in the tetrazolium dye. Cultured bacteria in log growth phase were centrifuged down to pellet, which was suspended using inoculating fluid (Biolog, Hayward, CA) to an OD value around 0.26. The suspension was added to the microplates and sealed by anaerobic GasPak EZ anaerobe gas pouch system with anaerobic indicators (BD, Franklin Lakes, NJ). The plates were read between 20-24hrs following inoculation with a pre-grown isolate using spectrophotometer microplate reader (Molecular Devices, LLC, San Jose, CA) and reading data acquisition was performed using SoftMax Pro 7 software (Molecular Devices, LLC, San Jose, CA). The procedure was performed in triplicate for each strain. The no carbon control reading was subtracted from each of the readings of the wells, and student’s t test was performed to test if the average reading was significantly different from zero.

### Comparative genomics analyses

A total of 79 *B. pseudolongum* genomes were included in the analyses, which included the four sequenced in this study and all 75 available *B. pseudolongum* genomes on GenBank (retrieved September 2021, **Supplemental Table 5A**). The pangenome was constructed using anvi’o vers 6.2 workflow [69, 70]. Briefly, this workflow 1) dereplicates genomes based on similarity score calculated using Sourmash vers 3.3 [71], 2) uses BLASTP to compute ANI identity between all pairs of genes, 3) uses the Markov Cluster Algorithm (MCL) [72] to generate homologous gene clusters (HGCs) based on all-versus-all sequence similarity, and 4) aligns amino acid sequences using MUSCLE [73] for each gene cluster. Each gene was assigned to core or accessory according to the hierarchical clustering of the gene clusters. Sourmash vers 3.3 [71] was used to compute ANI across genomes. Functional annotation of each secreted protein was performed employing the eggNOG database v5.0 [74] using eggNOG-mapper v2 [75] and the results were imported into the anvi’o contig database. Further functional annotation included PFAMs based on hidden Markov model (HMM) search to Pfam vers34.0 [76]. Protein-coding genes were also annotated to metabolic functionality categories using KEGG (Kyoto Encyclopedia of Genes and Genomes) [77]. GhostKOALA annotation tool [78] was used to assign KEGG Identifiers. Enrichment analyses were performed using Anvi’o pangenome pipeline that take COG functions across genomes and clade affiliation as the explanatory variable. The equality of proportions across clade affiliation was tested using a Rao score test, which generates an enrichment score as the test statistic and a p value. The q-value was then calculated from the p value to account for multiple testing using R package qvalue [79]. A COG function was considered enriched if the q-value was below 0.05.

The prediction of genes encoding extracellular enzymes possessing structurally related catalytic and carbohydrate-binding modules catalyzing hydrolysis, modification, or synthesis of glycoside bounds was performed using dbCAN2 [80] and dbCAN HMMdb (v.9) that was built using CAZy database (v.07302020) [38]. To identify signal-peptide specific sequence motifs, we employed the subcellular localization prediction tool PSORTb (v.3.0.2) [81].

### *In vitro* co-culture

Peritoneal macrophages (MΦ) and bone marrow derived dendritic cells (BMDCs) were isolated and then seeded onto 24 well plates in 1 ml RPMI complete medium as described [82]. Briefly, MΦ were collected 4 days after i.p. injection of Remel Thioglycollate solution (Thermo Fisher Scientific, Waltham, MA). BMDCs were generated from bone marrow cells treated with 10 ng/ml GM-CSF (R&D Systems, Minneapolis, MN) for 10 days. Loosely adherent immature BMDCs were collected and then CD11c+ DCs were enriched using CD11c positive selection kit (Stemcell Technologies, Cambridge, MA). Twenty-four hours after culture of purified subsets, the cells were stimulated with UV-killed *Bifidobacterium* bacteria or purified EPS for 24 hours. Bacterial cells were killed by UV exposure at 100 μJ/cm2 for four 15-minute cycles with a UV CrossLinker (Fisher Scientific, Hampton, NH). Culture supernatants were collected from whole bacteria or EPS stimulated cultures and ELISA for TNFα, IL-6, and IL-10 (BioLegend, San Diego, CA) performed. The myeloid cells from co-culture wells were collected and analyzed by flow cytometry.

### Flow cytometry

Cells were passed through 70-μm nylon mesh screens (Thermo Fisher Scientific, Waltham, MA) to produce single-cell suspensions. Cell suspensions were treated with anti-CD16/32 (clone 93, eBioscience) to block Fc receptors, and then stained for 30 minutes at 4°C with antibodies against surface molecules (**Supplementary Table 9**) and washed 2 times in FACS buffer [phosphate buffered saline (PBS) with 0.5% w/v Bovine serum albumin (BSA)]. Samples were analyzed with an LSR Fortessa Cell Analyzer (BD Biosciences), and data analyzed with FlowJo software version 10.6 (BD Biosciences).

### Immunohistochemistry

Peripheral LN, mesenteric LN, and small bowel segment (duodenal-jejunal junction) were excised and washed in cold PBS before freezing in OCT (Sakura Finetek, Torrance, CA) in histology blocks on dry ice and then stored at -80°C. LN cryosections were cut in triplicate at 5 μm using a Microm HM 550 cryostat (Thermo Fisher Scientific, Waltham, MA). Sections attached to slides were fixed with cold acetone/ethanol (1:1) solution and washed in PBS buffer (Lonza, Morristown, NJ). Primary antibodies and isotype controls (**Supplementary Table 9)** were added to slides for 1 hour in a humidified chamber. Sections were washed with PBS, blocked with 2.5% donkey serum and 2.5% goat serum, and incubated with secondary antibodies for 60 minutes. Slides were then fixed with 4% paraformaldehyde/PBS (Alfa Aesar, Haverhill, MA) for 5 minutes, incubated with 1% glycerol for 5 minutes, and Prolong Gold Antifade Mountant with or without DAPI (Thermo Fisher Scientific, Waltham, MA) was added before applying cover slips. Images were acquired using an Accu-Scope EXC-500 fluorescent microscope (Nikon, Melville, NY) and analyzed with Volocity image analysis software (PerkinElmer, Waltham, MA). The percentage positive staining area was quantified based on at least 2 independent experiments with 3 mice/group, 3 LNs/mouse, 3 sections/LN, and 3–5 fields/section.

### Mice experiments

Female C57BL/6 mice between 8 and 14 weeks of age were purchased from The Jackson Laboratory (Bar Harbor, ME). All the procedures involving mice were performed in accordance with the guidelines and regulations set by the Office of Animal Welfare Assurance of the University of Maryland School of Medicine, under the approved IACUC protocols 0518004 and 0121001. Mice were fed antibiotics (kanamycin, gentamicin, colistin, metronidazole, and vancomycin) *ad libitum* in drinking water on days -6 to -1. On day 0, cultured *Bifidobacterium* ATCC25526 or UMB-MBP-01 were gavaged p.o. Mice received tacrolimus (3 mg/kg/d s.c.) on days 0 and 1. On day 2, the animals were euthanized. Mesenteric and peripheral (axillary, inguinal, popliteal, brachial) LNs as well as small intestine were harvested. Fecal pellets were also collected prior to euthanasia into individual tubes, using aseptic technique to minimize handling, and stored at -80°C. Antibiotics were USP grade or pharmaceutical secondary standard (all from MilliporeSigma): kanamycin sulfate (0.4 mg/ml), gentamicin sulfate (0.035 mg/ml), colistin sulfate (850 U/ml), metronidazole (0.215 mg/ml), and vancomycin hydrochloride (0.045 mg/ml) were dissolved in vivarium drinking water and administered *ad libitum*. Tacrolimus (USP grade, MilliporeSigma) was reconstituted in DMSO (USP grade, MilliporeSigma) at 20 mg/ml and diluted with absolute ethanol (USP grade, Decon Labs, King of Prussia, PA) to 1.5mg/ml. DMSO/ethanol stock was diluted 1:5 in sterile PBS for s.c. injection and injected at 10 μl/g (3 mg/kg/day).

### Metagenomic sequencing and microbiome analyses

Harvested intestine tissues and luminal contents were stored immediately in DNA/RNA Shields (Zymo Research, Irvine, CA) to stabilize and protect the integrity of nucleic acids and minimize the need to immediately process or freeze specimens. The colon content from ∼1cm colon tissue was used in DNA extraction. Metagenomic sequencing libraries were constructed from the same DNA using the Nextera XT Flex kit (Illumina) according to the manufacturer recommendations. Libraries were then pooled together in equimolar proportions and sequenced on a single Illumina NovaSeq 6000 S2 flow cell at the Genomic Resource Center of the Institute for Genome Sciences at the University of Maryland School of Medicine.

Metagenomic sequence reads were removed using BMTagger v3.101 [83] using a Genome Reference Consortium Mouse Build 39 of strain C57BL/6J (GRCm39) [84]. Sequence read pairs were removed even if only one of the reads matched to the mice genome reference. The Illumina adapter was trimmed and quality assessment was performed using default parameters in fastp (v.0.21.0) [85]. The taxonomic composition of the microbiomes was established using Kraken2 (v.2020.12) [86] and Braken (v. 2.5.0) [87] using the comprehensive mouse gut metagenome catalog (CMGM) [47] to calculate the metagenomic taxonomic composition. The Phyloseq (v.1.34.0) [88] R package was used to generate the diversity plot and barplot. Linear discriminant analysis (LDA) effect size (LEfSe) analysis [48] was used to identify fecal phylotypes that could explain differences between. For LEfSe, only taxonomic groups present in >1% of at least one sample were included in the analyses; the alpha value for the non-parametric factorial Kruskal-Wallis (KW) sum-rank test was set at 0.05 and the threshold for the logarithmic LDA model [89] score for discriminative features was set at 2.0. An all-against-all BLAST search in multi-class analysis was performed. Metagenomics dataset was mapped to the protein database UniRef90 [90] to ensure the comprehensiveness in functional annotation, and was then summarized using HUMAnN2 (Human Microbiome Project Unified Metabolic Analysis Network) (v0.11.2) [91] to efficiently and accurately determine the presence, absence, and abundance of metabolic pathways in a microbial community. Further, HUMAnN2 employed a tiered search strategy enabling a species-resolved functional profiling of metagenomes, hence to characterize the contribution to the functional pathways of a known species. Canonical Correspondence Analysis (CCA) was used in ordination analysis, and biplot was generated using vegan package [92, 93] based on bray-curtis distance. CA1 and CA2 were selected as the major components based on the eigenvalue. A species score was scaled proportional to the eigenvalues representing the direction from the origin where the group had a larger than average value for the particular species [92, 94]. The species scores greater than 1 were used to select the species that were considered the most significant contributors to each group.

### RNA isolation, transcriptome sequencing and analyses of the intestinal tissues

Dissected intestinal tissues were stored immediately in RNAlater solution (QIGEN) that had been stored at −80°C to stabilize and protect the integrity of RNA [95]. For each sample, bulk RNA was extracted from ∼1cm of intestine tissues. Prior to the extraction, 500µl of ice-cold RNase free PBS was added to the sample. To remove the RNAlater, the mixture was centrifuged at 8,000x*g* for 10min and the resulting pellet resuspended in 500µl ice-cold RNase-free PBS with 10µl of β-mercaptoethanol. Tissue suspension was obtained by bead beating procedure using the FastPrep lysing matrix B protocol (MP Biomedicals, Solon, OH) to homogenized tissues. RNA was extracted from the resulting suspension using TRIzol Reagent (Invitrogen, Carlsbad, CA) following the manufacturer recommendations and followed by protein cleanup using Phasemaker tubes (Invitrogen) and precipitation of total nucleic acids using isopropanol. RNA was resuspended in DEPC-treated DNAase/RNAase-free water. Residual DNA was purged from total RNA extract by treating once with TURBO DNase (Ambion, Austin, TX, Cat. No. AM1907) according to the manufacturer’s protocol. DNA removal was confirmed via PCR assay using 16S rRNA primer 27F (5′-AGAGTTTGATCCTGGCTCAG-3′) and 534R (5′-CATTACCGCGGCTGCTGG-3′). The quality of extracted RNA was verified using the Agilent 2100 Expert Bioanalyzer using the RNA 1000 Nano kit (Agilent Technologies, Santa Clara, CA). Ribosomal RNA depletion and library construction were performed using the RiboZero Plus kit and TruSeq stranded mRNA library preparation kit (Illumina) according to the manufacturer’s recommendations. Libraries were then pooled together in equimolar proportions and sequenced on a single Illumina NovaSeq 6000 S2 flow cell at the Genomic Resource Center (Institute for Genome Sciences, University of Maryland School of Medicine) using the 150 bp paired-end protocol.

Bioinformatic analysis of the transcriptome data includes the quality of fastq files, which was evaluated by FastQC. Reads were aligned to the mouse genome (v. Mus_musculus.GRCm39) using HiSat (v. HISAT2-2.1.0) [96] and the number of reads that aligned to the coding regions were determined using HTSeq (v.1.0.0) [97].

Significant differential expression was assessed using DEseq with an FDR value ≤ 0.05 [98]. Quadrant plot to show whether DEGs have the same or opposite relationships between each of the pairwise comparisons of UMB-MBP-01 vs ATCC25526 and UMB-MBP-01 vs control. Gene Ontology (GO) enrichment analysis was performed in order to identify GO terms significantly over-represented in genes deregulated in specific comparisons and, as a result, to suggest possible functional characteristics of these genes. Enriched GO terms in the set of genes that are significantly over-expressed or under-expressed in a specific condition may suggest possible mechanisms of regulation or functional pathways that are, respectively, activated or repressed in that condition. Over-representation analyses [99] of differentially expressed genes (DEGs) against GO ontologies was performed using enrichGO function of clusterProfile Bioconductor package [100]. Cnetplot function was used to depict the linkages of genes and GO terms as a Gene-Concept Network for top over-represented GO terms based on q-value and gene-count.

## Data Availability

The assembly of the genomes (accession numbers were listed in **Supplemental Table 2**), metagenome (SRP361281), and transcriptome sequences were submitted to GenBank under BioProject PRJNA809764.

## Contributions

B. M., E. F. M, J. B. designed the experiments. S. J. G., V. S., W. P., R. L., L. L., C. P., M.W. conducted and analyzed the *in vitro* and *in vivo* experiments. H. W. L. performed the AN Biolog assay and RNA extraction. L. H., E. F. M., A. M. performed colony isolation and strain characterization. B. M. and Y. S. performed the bioinformatics analyses. B. M., M. F. performed the statistical analyses. B. M., S. J. G., M. F., E. F. M., and J. S. B. wrote the manuscript.

## Acknowledgements

This work was supported by National Institute of Health (NIH) National Heart, Lung and Blood Institute award R01HL148672 (JSB/BM) and NIH National Institute of Allergy and Infectious Diseases training grant T32AI95190-10 (SJG). This manuscript has been released as a pre-print at BioRxiv XX. The authors thank C. Colin Brinkman and Yanbao Xiong at the University of Maryland for helping with pilot data and initial experiment preparation. Emmanuel Mongodin, PhD, contributed to this work as an employee of the University of Maryland School of Medicine. The views expressed in this manuscript are his own and do not necessarily represent the views of the National Institutes of Health or the United States Government.

