## Supplemental_Materials for "Strain-specific alterations in gut microbiome and host immune responses elicited by *Bifidobacterium pseudolongum*"

**Supplemental Figures**

**Supplemental Figure 1.** Proportion of the pangenome categories of *Bifidobacterium* species: *B. breve*, *B. longum* and *B. pseudolongum*. Homologous gene clusters (HGCs) identified based on all-versus-all sequence similarity. Each gene was assigned to core or accessory according to the hierarchical clustering of the gene clusters.

**Supplemental Figure 2.** Glycosyl hydrolases similarity among *B. pseudolongum* genomes. BLAST Score Ratio (BSR) approach [1] with default settings was used to calculate the GH similarity. UMB-MBP-01 was used as the reference to compare to glycosyl hydrolyse capabilities of the other 61 *B. pseudolongum* strains. Color indicates BSR score scaled from 0 (absent) to 100 (identical proteins).

**Supplemental Figure 3.** Flow cytometry gating strategy.

**Supplemental Figure 4.** C57BL/6 mice treated with antibiotics for 6 days and then gavaged with PBS, ATCC25526 (ATCC) or UMB-MBP-01 (MD) UV-killed *Bifidobacteria* or EPS derived from each strain. Mice then treated with tacrolimus (3 mg/kg/d s.c.) for the next two days. Tissues harvested 2 days after bacterial gavage and analyzed by flow cytometry. MLN stained for **A**) B220, **B**) CD4, **C**) CD8, **D**) CD4 and Foxp3, **E**) CD11c, and **F**) F4/80. PLN stained for **G**) B220, **H**) CD4, **I**) CD8, **J**) CD4 and Foxp3, **K**) CD11c, and **L**) F4/80. Frozen tissue sections analyzed by immunohistochemistry for **M**) CD11c and **N**) F4/80. **O**) MLN stained for laminin $\alpha 4$ and $\alpha 5$, with their ratio depicted. Treatment groups compared using one-way ANOVA. ** p value < 0.01. Data representative of two separate experiments, 3 mice per group.

**Supplemental Figure 5.** Gene-Concept network for most over-represented Gene Ontology (GO) terms depict over-represented functions based on q-value and gene-count. Over-representation analyses [2] of differential expressed genes (DEGs) that are only differentially abundant in **A**) UMB-MBP-001 vs control up-regulated; **B**) UMB-MBP-001 vs control down-regulated; **C**) UMB-MBP-001 vs ATCC25526 up-regulated; **D**) UMB-MBP-001 vs ATCC25526 down-regulated; **E**) ATCC25526 vs control up-regulated; **F**) ATCC25526 vs control down-regulated, using GO ontologies performed using enrichGO function of clusterProfile Bioconductor package[3]. DEGs were determined using log2 fold change (LFC) >(+/-)1 and false discovery rate (FDR)<0.05.

**Supplemental Figure 6.** Transcriptome profiling of intestinal tissues in response to live *Bifidobacterium* cells or *Bifidobacterium* exopolysaccharides (EPS). **A**) MA plot of *Bifidobacterium* group vs the no bacteria control, DEGs of B cell activation and signaling pathways labeled. **B**) Quadrant plot showing whether DEGs have the same or opposite relationships between each of the pairwise comparisons of *Bifidobacterium* vs control and EPS vs control. Each point represents a DEG. Gene-pathway network for the top 5 over-represented GO terms in comparison of **C**) *Bifidobacterium* live vs no bacteria control and **D**) EPS vs no bacteria control. Raw counts generated by HTSeq used for differential expression (DE) analyses using DESeq2 [4] to estimate gene dispersion, normalize counts by library size and statistically determine DE genes (DEGs) using log2 fold change (LFC) >(+/-)1 and false discovery rate (FDR)<0.05. Over-representation analyses of DEGs using GO ontologies were performed using enrichGO function of clusterProfile Bioconductor package [3]. Cnetplot function used to depict the linkages of genes and GO terms as a Gene-Concept Network for over-represented GO terms based on q-value and gene-count.

**Supplemental Figure 7.** Comparison of relative abundance of bacterial groups in stool samples of the mice in different experiment groups. Cumulative abundance of bacteria groups from the four most abundant phyla present in all mice shown.

**Supplemental Figure 8**. Taxonomic groups differentially abundant with statistical and biological significance, ranked according to the effect size. Only taxonomic groups present at >1% in at least one sample were included in the analyses. Logarithmic linear discriminant analysis (LDA) effect size (LEfSe) was employed (Segata et al., 2011). The alpha threshold value for the pairwise non-parametric Kruskal-Wallis test was 0.05 and the threshold for the logarithmic LDA model score (Fisher, 1936) for discriminative features was 2.0. All-against-all comparison in multi-class analysis were performed.

**Supplemental Figure 9.** PCA analysis between ATCC25526 and UMB-MBP-01 groups. Canonical Correspondence Analysis (CCA) was used in ordination analysis of **A**) taxonomic profiles and **B**) functional profiles. Biplot was generated using vegan package [5, 6] based on Bray-Curtis distance. CA1 and CA2 were selected as the major components based on the eigenvalue. Microbial functional pathways were characterized using HUMAnN2 (v0.11.2)[7] and Uniref90 database [8].

**Supplemental Tables:**

**Supplemental Table 1.** Statistics of pangenome analyses. **A**) Statistics of the 79 *Bifidobacterium pseudolongum* strains including the 4 strains sequenced in this study; **B**) homologous gene clusters (HGCs) of *Bifidobacterium pseudolongum* generated in metapangenome analyses; **C**) Comparison of protein coding genes among the four bifidobacterial strains from the same source; **D**) Comparison of protein coding genes between UMB-MBP-01 and ATCC25526; **E**) HGCs enriched in different clades; **F**) Comparison of GHs (glycoside hydrolyses) between UMB-MBP-01 and ATCC25526.

**Supplemental Table 2.** Information of the genomic and metagenomes sequencing. **A**) Statistics of genomic sequencing reads and assembly; **B**) Statistics of the metagenomes sequencing of intraluminal stools.

**Supplemental Table 3.** Anaerobic microplates pre-coated with various carbon sources the carbohydrate utilization capabilities of UMB-MBP-01 and ATCC25526.

**Supplemental Table 4.** Information on glycosyl hydrolases (GH) characterization of *B. pseudolongum* pangenome using Carbohydrate-Active enZYmes Database (CAZy) database [9]. **A**) GHs of the HGCs generated in pangenome analyses; **B**) GH similarity among *B. pseudolongum* genomes. UMB-MBP-01 was used as the reference to compare to glycohydrolyses capabilities of all other 61 *B. pseudolongum* strains. BLAST Score Ratio (BSR) approach [1] in default setting used to calculate similarity score.

**Supplemental Table 5.** Protein localization of the coding genes based on the presence of a signal peptide [10]. **A**) localization information of all homologous gene clusters (HGCs) of *Bifidobacterium pseudolongum*; **B**) Distribution of all *B. pseudolongum* HGCs localization categories; **C**) proteins predicted to be associated with cell wall; **D**) proteins predicted to be extracellular.

**Supplemental Table 6.** Differential expressed genes (DEGs) between each of the pairwise comparison of UMB-MBP-001 vs control and UMB-MBP-001 vs ATCC25526. **A**) List of DEGs according to bin name listed in the table correspond to the quadrant plot in **Figure 5A**; **B**) DEGs between the comparison of UMB-MBP-001 vs control; **C**) DEGs between the comparison of ATCC25526 vs control; **D**) DEGs between the comparison of UMB-MBP-001 vs ATCC25526.

**Supplemental Table 7.** Taxonomic profiles of the metagenomes sequencing of intraluminal stools. Taxonomy was characterized using the comprehensive mouse gut metagenome catalog (CMGM) [11] designed specifically to characterize mouse gut microbiome.

**Supplemental Table 8.** Statistics of Canonical Correspondence Analysis (CCA). CA1 and CA2 are selected as the major components based on the eigenvalue. CCA was generated using vegan package [5, 6] based on bray-curtis distance.

**Supplemental Table 9.** Primary and secondary antibodies used in this study.

**SFigure 1.**

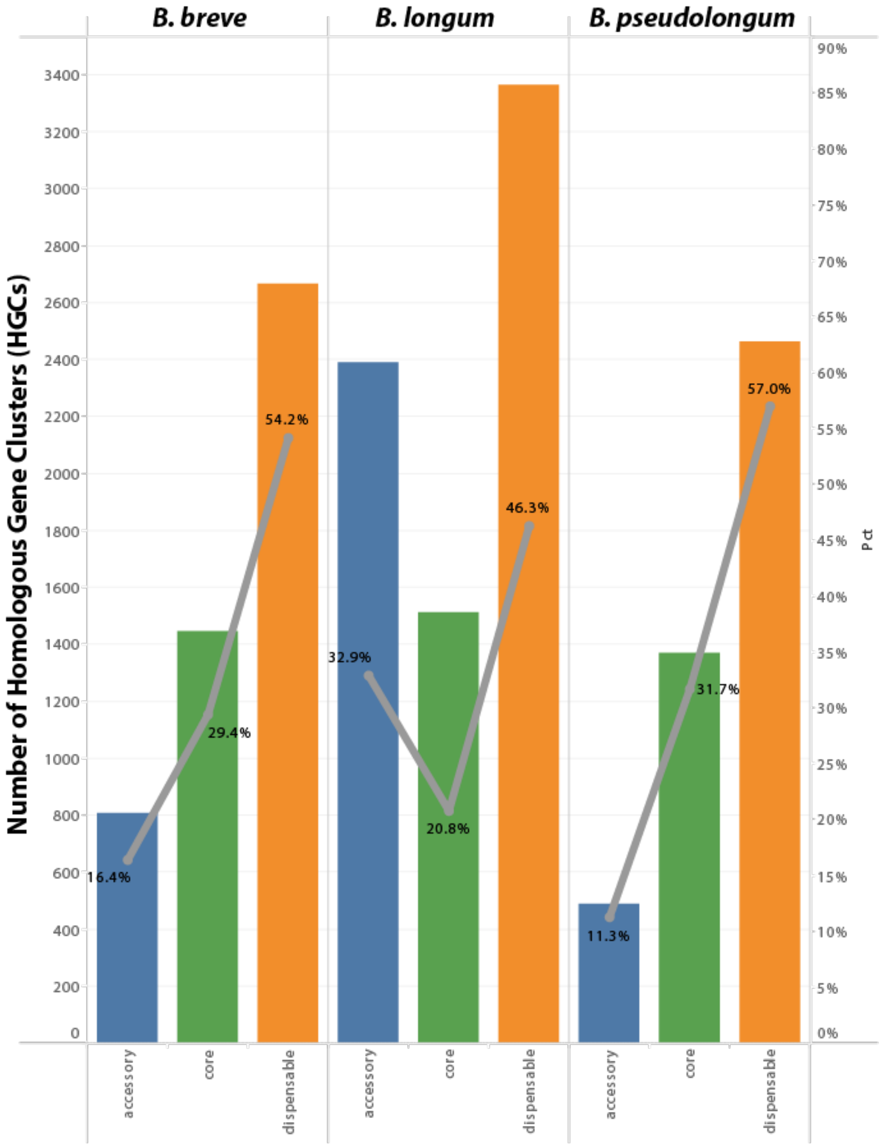


| *B breve* | count | % | *B longum* | count | % | *B pseulongum* | count | % |
|---|---|---|---|---|---|---|---|---|
| accessory | 808 | 16.4% | accessory | 2,391 | 32.9% | accessory | 488 | 11.3% |
| core | 1,448 | 29.4% | core | 1,511 | 20.8% | core | 1,370 | 31.7% |
| dispensable | 2,666 | 54.2% | dispensable | 3,363 | 46.3% | dispensable | 2,463 | 57.0% |
| total | 4,922 | | total | 7,265 | | total | 4,321 | |

**SFigure 2.**

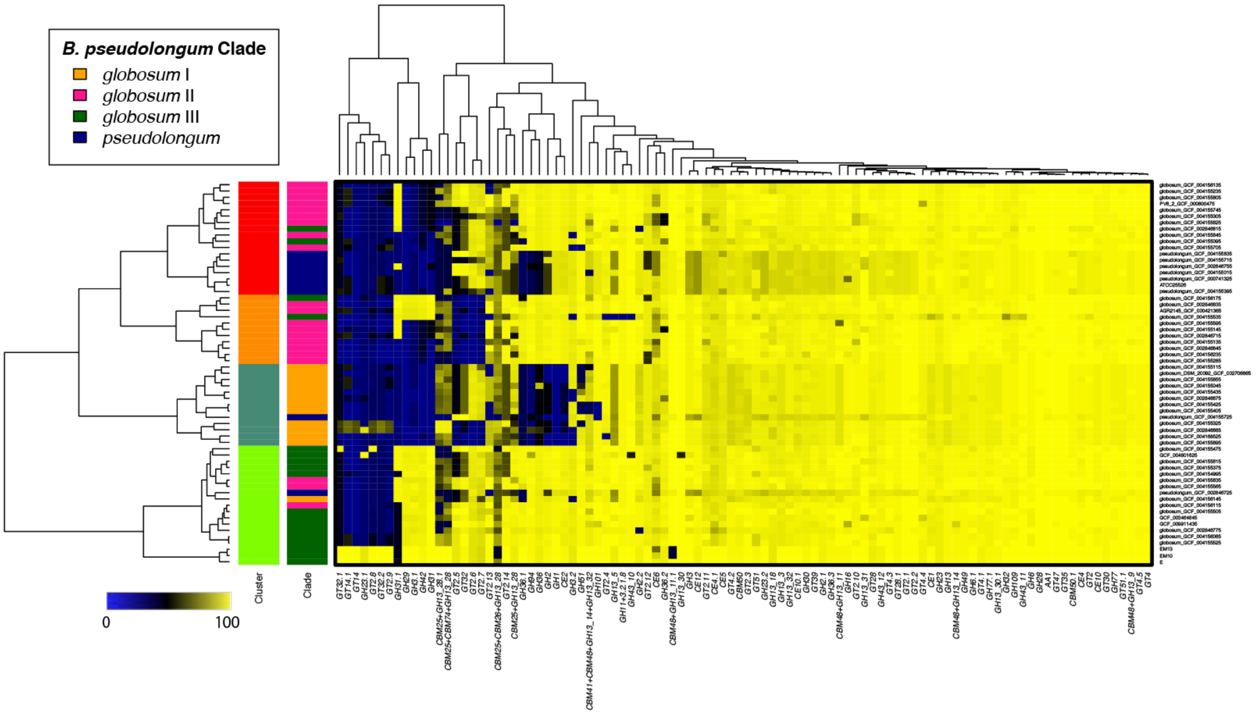
B. pseudolongum Clade
globosum I
globosum II
globosum III
pseudolongum
Cluster
Clade
0
100

**SFigure 3.**

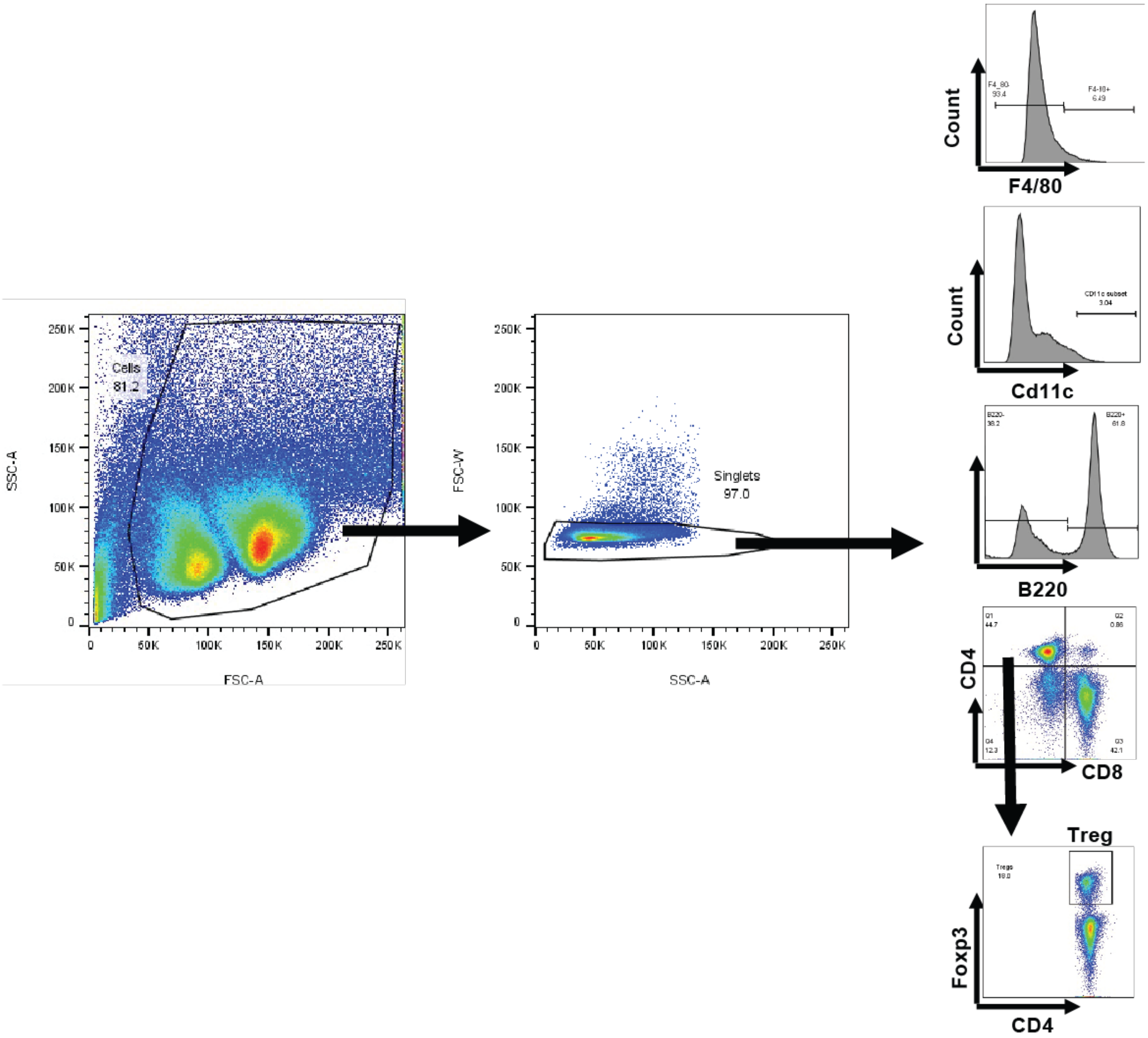

Cells
81.2
SSC-A
FSC-A
Singlets
97.0
FSC-W
SSC-A
Count
F4/80
Count
Cd11c
B220
CD4
CD8
Treg
Foxp3
CD4

SFigure 4.

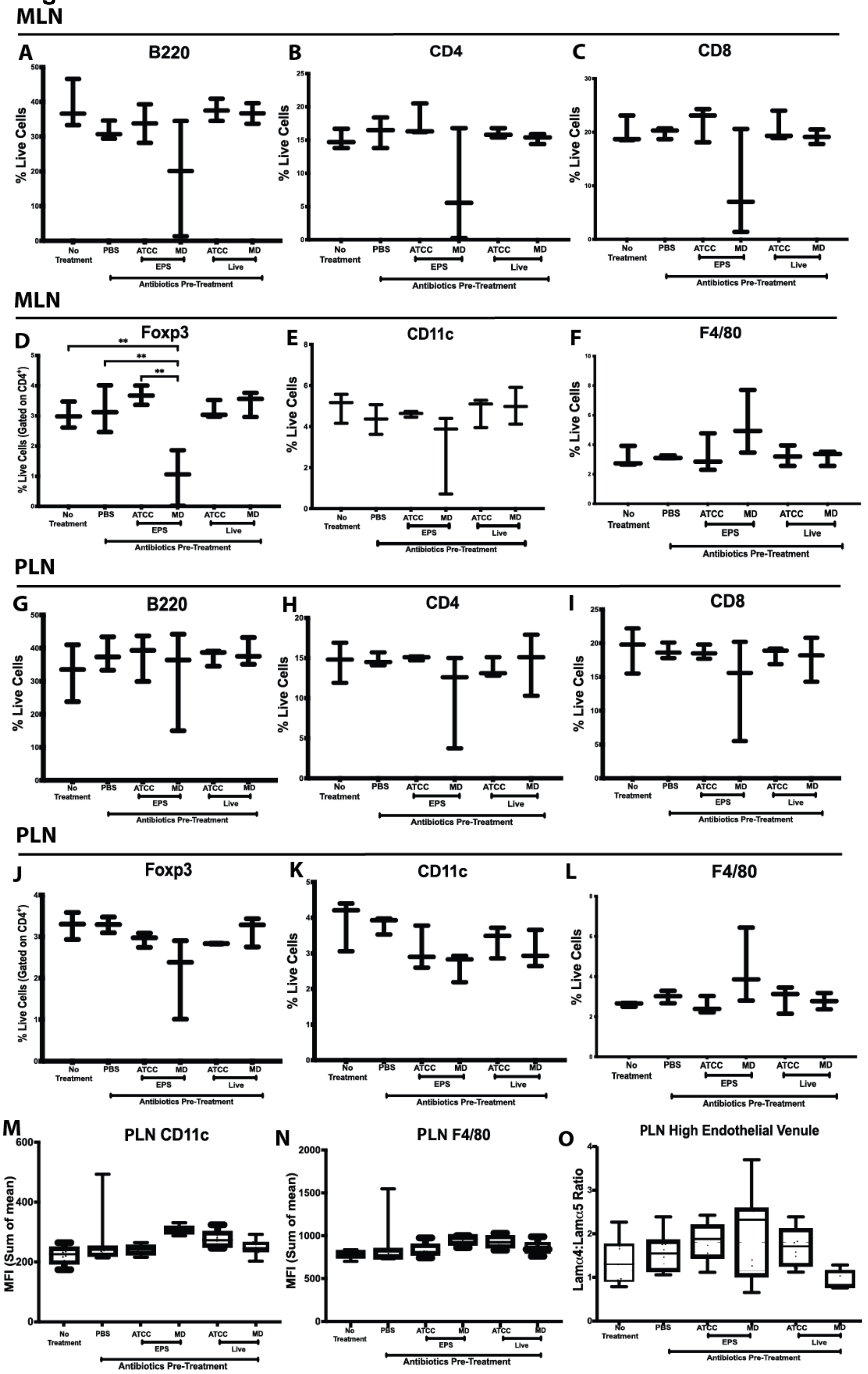
MLN
A
B220
% Live Cells
No Treatment
PBS
ATCC
MD
EPS
Live
Antibiotics Pre-Treatment
B
CD4
C
CD8
D
Foxp3
**
% Live Cells (Gated on CD4+)
E
CD11c
F
F4/80
PLN
G
B220
H
CD4
I
CD8
J
Foxp3
K
CD11c
L
F4/80
M
PLN CD11c
MFI (Sum of mean)
N
PLN F4/80
O
PLN High Endothelial Venule
Lamα4:Lamα5 Ratio

**SFigure 5.**

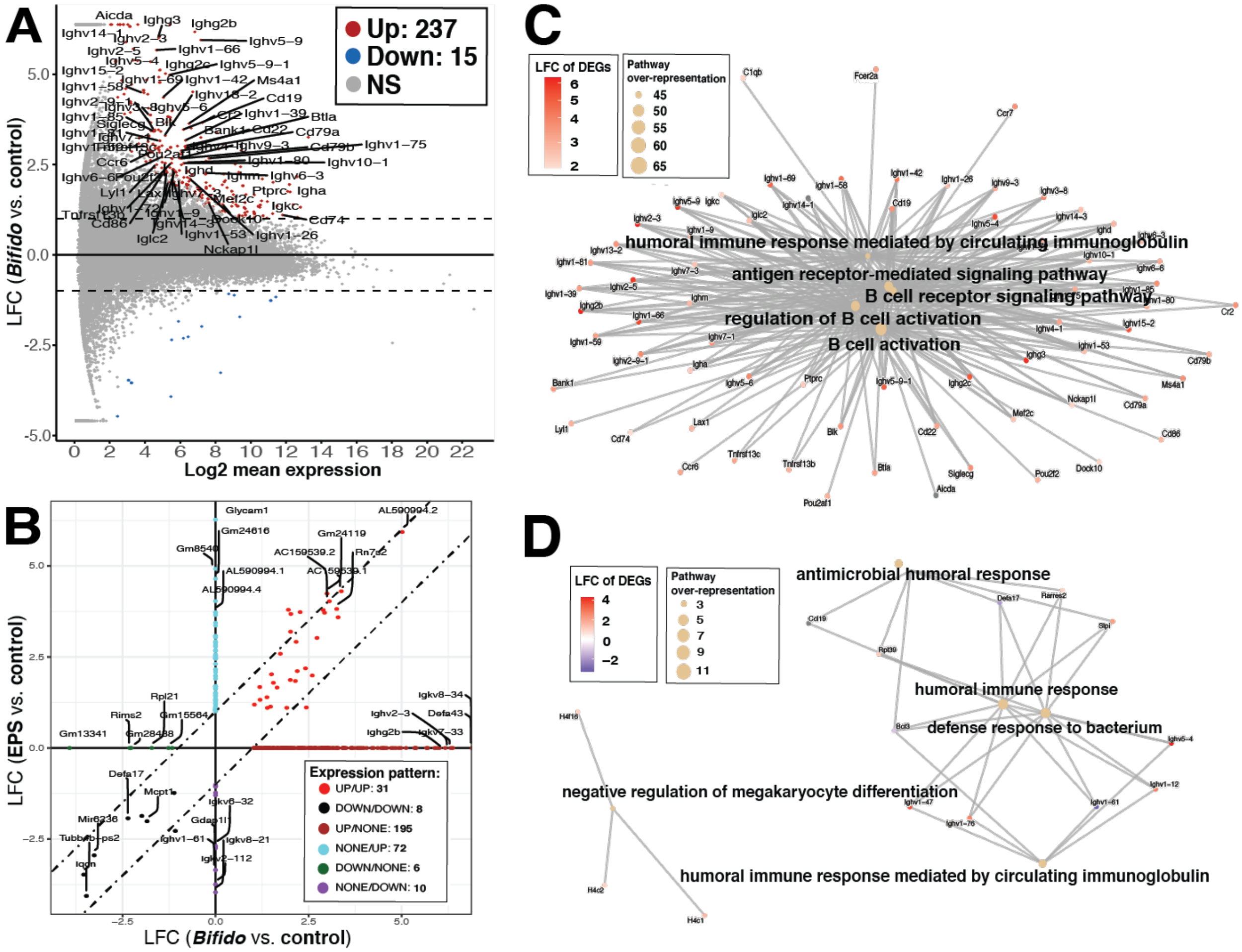
A
Up: 237
Down: 15
NS
LFC (Bifido vs. control)
Log2 mean expression
B
LFC (EPS vs. control)
LFC (Bifido vs. control)
Expression pattern:
UP/UP: 31
DOWN/DOWN: 8
UP/NONE: 195
NONE/UP: 72
DOWN/NONE: 6
NONE/DOWN: 10
C
LFC of DEGs
Pathway over-representation
humoral immune response mediated by circulating immunoglobulin
antigen receptor-mediated signaling pathway
B cell receptor signaling pathway
regulation of B cell activation
B cell activation
D
antimicrobial humoral response
humoral immune response
defense response to bacterium
negative regulation of megakaryocyte differentiation
humoral immune response mediated by circulating immunoglobulin

### SFigure 6.

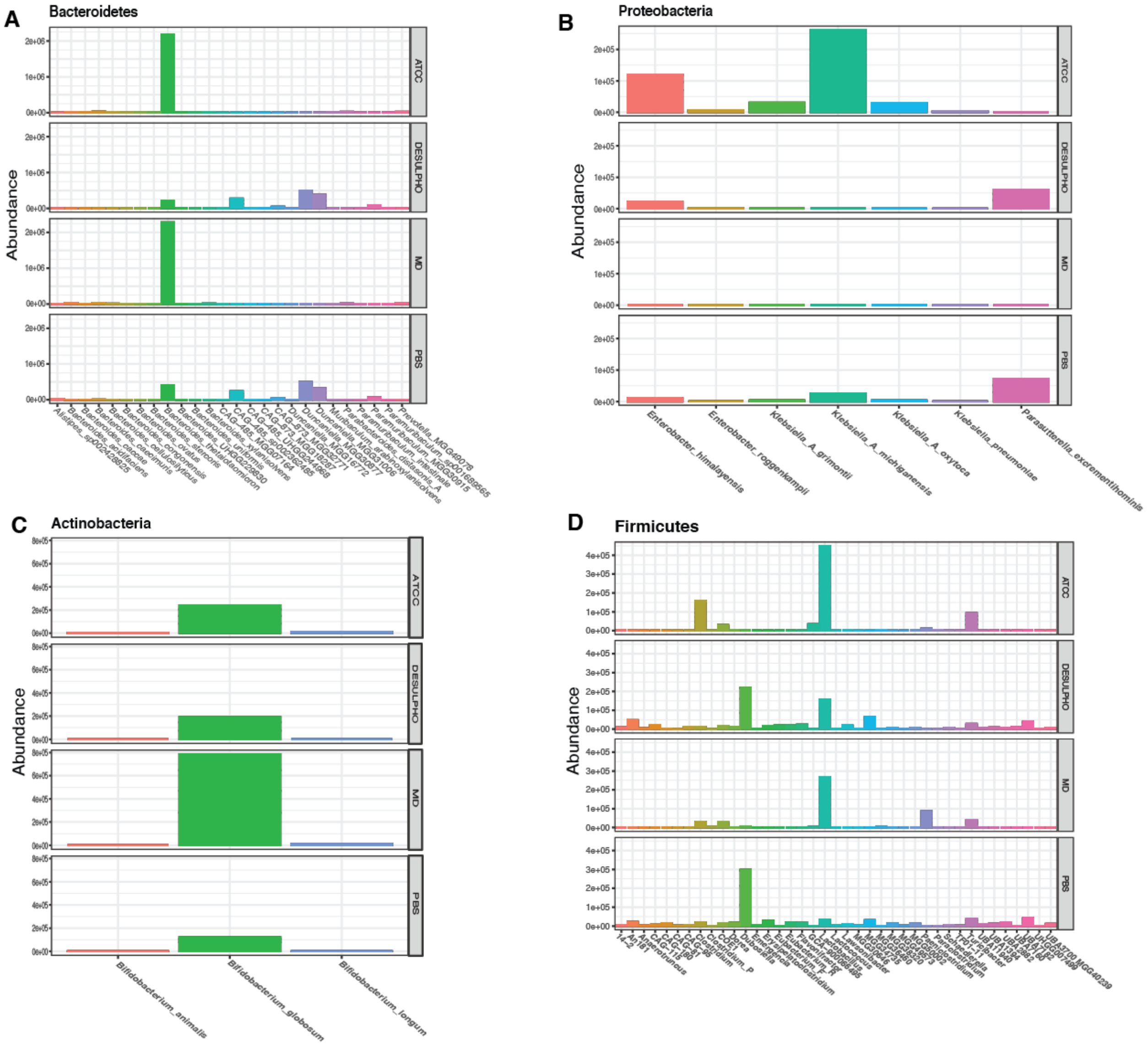
A
Bacteroidetes
B
Proteobacteria
C
Actinobacteria
D
Firmicutes
Abundance
ATCC
DESULPHO
MD
PBS
Enterobacter_himalayensis
Enterobacter_roggenkampii
Klebsiella_A_grimontii
Klebsiella_A_michiganensis
Klebsiella_A_oxytoca
Klebsiella_pneumoniae
Parasutterella_excrementihominis
Bifidobacterium_animalis
Bifidobacterium_globosum
Bifidobacterium_longum

**SFigure 7.**

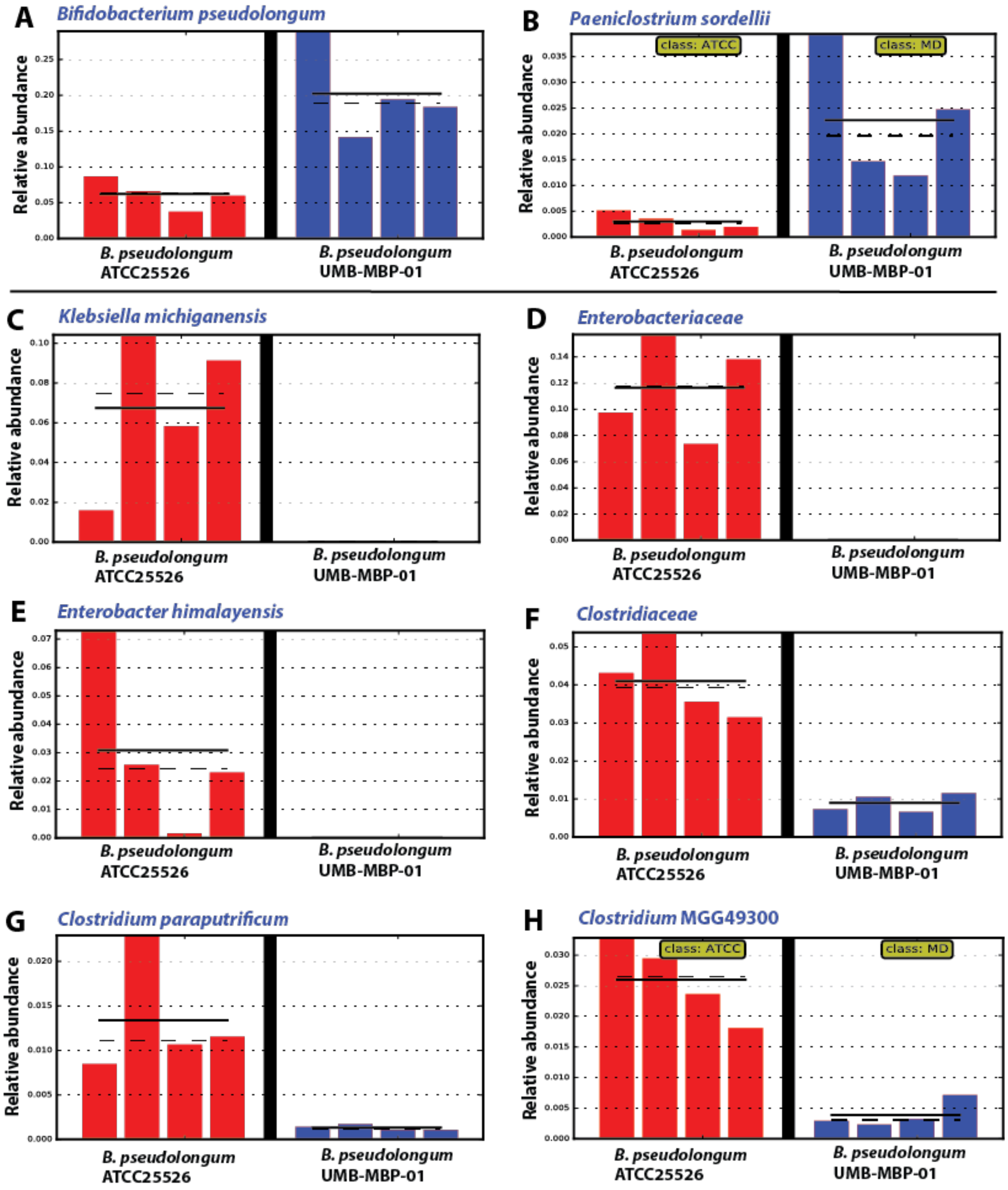
A
Bifidobacterium pseudolongum
B
Paeniclostrium sordellii
class: ATCC
class: MD
C
Klebsiella michiganensis
D
Enterobacteriaceae
E
Enterobacter himalayensis
F
Clostridiaceae
G
Clostridium paraputrificum
H
Clostridium MGG49300
Relative abundance
B. pseudolongum
ATCC25526
B. pseudolongum
UMB-MBP-01

**SFigure 8.**

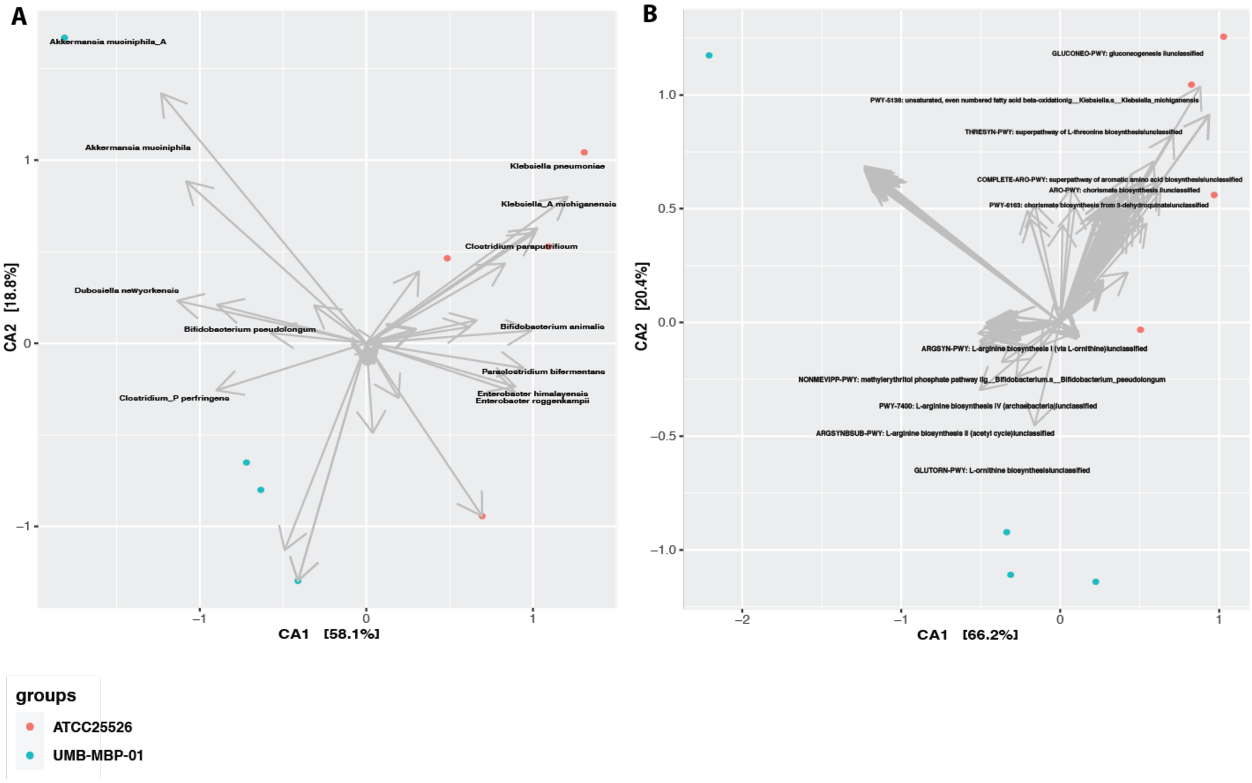
A
Akkermansia muciniphila_A
Akkermansia muciniphila
Klebsiella pneumoniae
Klebsiella_A michiganensis
Clostridium paraputrificum
Dubosiella newyorkensis
Bifidobacterium pseudolongum
Bifidobacterium animalis
Paraclostridium bifermentans
Clostridium_P perfringens
Enterobacter himalayensis
Enterobacter roggenkampii
CA1 [58.1%]
CA2 [18.8%]
B
GLUCONEO-PWY: gluconeogenesis I|unclassified
PWY-5138: unsaturated, even numbered fatty acid beta-oxidation|g__Klebsiella.s__Klebsiella_michiganensis
THRESYN-PWY: superpathway of L-threonine biosynthesis|unclassified
COMPLETE-ARO-PWY: superpathway of aromatic amino acid biosynthesis|unclassified
ARO-PWY: chorismate biosynthesis I|unclassified
PWY-6163: chorismate biosynthesis from 3-dehydroquinate|unclassified
ARGSYN-PWY: L-arginine biosynthesis I (via L-ornithine)|unclassified
NONMEVIPP-PWY: methylerythritol phosphate pathway I|g__Bifidobacterium.s__Bifidobacterium_pseudolongum
PWY-7400: L-arginine biosynthesis IV (archaebacteria)|unclassified
ARGSYNBSUB-PWY: L-arginine biosynthesis II (acetyl cycle)|unclassified
GLUTORN-PWY: L-ornithine biosynthesis|unclassified
CA1 [66.2%]
CA2 [20.4%]
groups
ATCC25526
UMB-MBP-01

### SFigure 9.

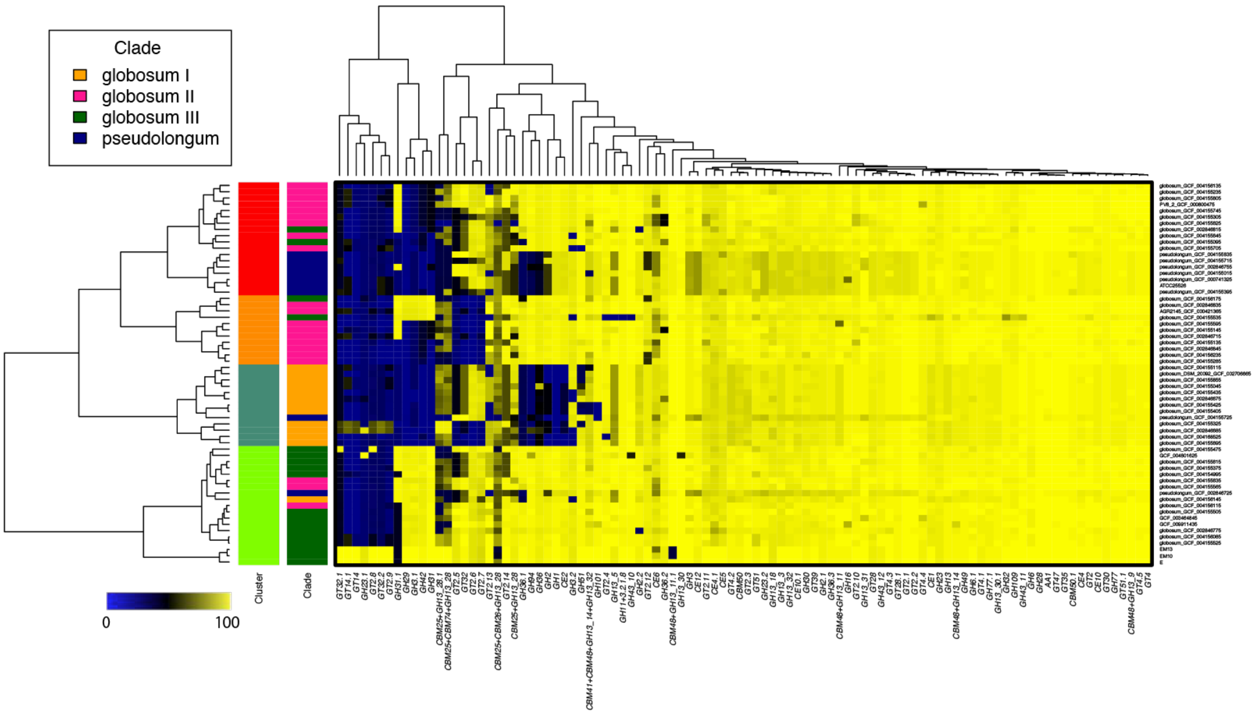
Clade
globosum I
globosum II
globosum III
pseudolongum
Cluster
Clade
0
100
